# Using NAMs to characterize chemical bioactivity at the transcriptomic, proteomic and phosphoproteomic levels

**DOI:** 10.1101/2022.05.18.492410

**Authors:** Yuan Li, Zhenpeng Zhang, Songhao Jiang, Feng Xu, Liz Tulum, Kaixuan Li, Shu Liu, Suzhen Li, Lei Chang, Mark Liddell, Fengjuan Tu, Xuelan Gu, Paul Lawford Carmichael, Andrew White, Shuangqing Peng, Qiang Zhang, Jin Li, Tao Zuo, Predrag Kukic, Ping Xu

## Abstract

Omic-based technologies are of particular interest and importance for non-animal chemical hazard and risk characterization based on the premise that any apical endpoint change must be underpinned by some alterations measured at the omic levels. In this work we studied cellular responses to caffeine and coumarin by generating and integrating multi-omic data from transcriptomic, proteomic and phosphoproteomic experiments. We have shown that the methodology presented here is able to capture the complete chain of events from the first compound-induced changes at the phosphoproteome level to changes in gene expression induced by transcription factors and lastly to changes in protein abundance that further influence changes at the cellular level. In HepG2 cells we found the metabolism of lipids and general cellular stress to be dominant biological processes in response to caffeine and coumarin exposure, respectively. The phosphoproteomic changes were detected early in time, at very low concentrations and provided a fast adaptive cellular response to chemical exposure. Changes in protein abundance were found much less frequently than the transcriptomic changes and can be used, together with the transcriptomic changes, to facilitate a more complete understanding of pathway responses to chemical exposure.

**GRAPHIC ABSTRACT:** 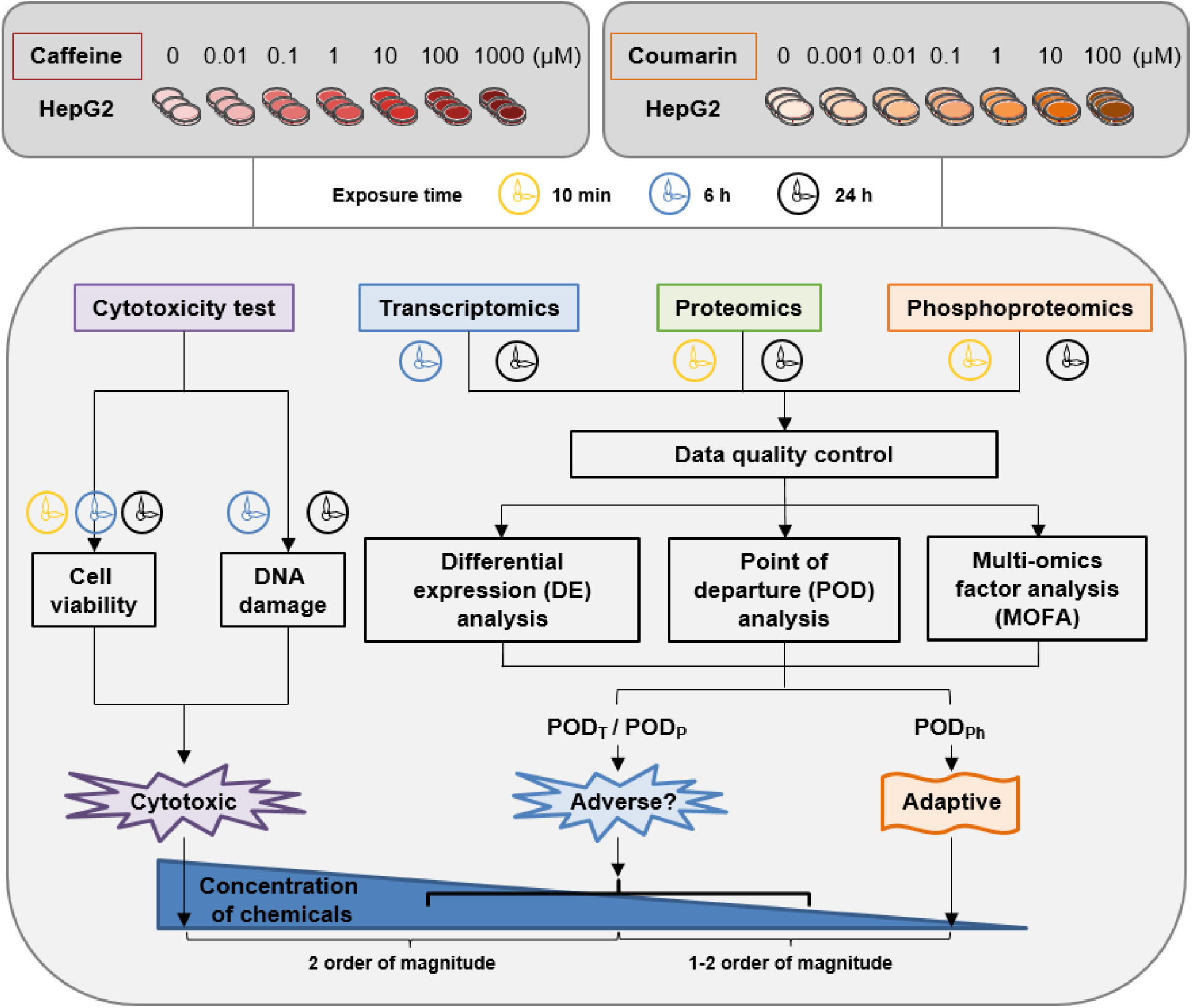

## INTRODUCTION

In 2018, the U.S. Environmental Protection Agency (EPA) proposed the concept of new approach methodologies (NAMs), which were defined as any technology or methodology mainly focused on effectively providing information on chemical hazard and risk without using the whole animal models^1^. This broad concept includes any existing or new *in vitro* bioactivity studies, *in silico* modelling and cheminformatics approaches and combinations thereof that inform on the bioactivity and exposure of the chemical. Among existing NAMs, the omic-based technologies are of particular importance based on the assumption that any apical endpoint change indicative of impaired health must be underpinned by some alterations at the omic level, including the transcriptome, metabolome, proteome, epigenome and genome^2–4^.

High throughput transcriptomics (HTTr) has already found its application in chemical bioactivity characterization due in a large part to its relatively low cost, high S/N ratio, need for a small sample size and most importantly broad biological coverage^5–8^. HTTr uses RNA-seq-based multiplexed read-outs of gene expression to measure mRNA changes directly from cell lysates in 384-well format and hence allows for cost-efficient screening of a large number of chemicals in a concentration-response manner^9^. Accumulating studies have revealed that point of departure (POD), as defined by the most sensitive transcriptomics gene, pathway or gene signature alterations in *in vitro* cellar assays, appear to be conservative or within similar order of magnitude to the doses causing pathological outcomes in long-term animal studies^5, 10^. Despite their protective nature, mechanistically speaking, it is not completely clear how and if HTTr-determined PODs are linked to the functional adversity in the cell or tissue caused by the chemical exposure. This may be due to a lack of our current biological understanding of how the changes are linked to adversity, lack of correlation between transcript and protein levels, or due to the limitations in the temporal nature of the assay. Nevertheless, it remains a source of uncertainty that could result in over-conservatism when using PODs determined from HTTr studies for chemical safety decision making.

The use of other omic technologies may provide additional or confirmative biomarkers to complement and reduce the uncertainty in HTTr assay-based decision-making and further build confidence in using NAM-based omic technologies for risk assessment. In recent years, technological progress in proteomics has offered quantitative analysis of tissues and cells encompassing the entire proteome^11^. Notably, it is now possible to determine the relative abundance of proteins in samples directly from proteomics data and without laborious preparation of standards or generation of calibration curves^12^. From the functional perspective, proteins are products of gene expression that mediate biochemical activities of cells and tissues and therefore are closer to the functional level. Besides the transcriptionally controlled effects on protein abundance, there are translational and also posttranslational effects that affect protein synthesis and stability. Indeed, ample evidence has shown RNA expression levels for a specific transcript product correlate poorly to its protein expression level^13–16^. Consequently, determination of differentially expressed proteins in the cell would help understand possible implications of measured transcriptomic changes and build confidence in the detection of pathway responses to chemical exposure.

The use of proteomics and phosphoproteomics along with other omics has been explored lately for applications in toxicology. Most notably, the CEFIC LRI-funded project called “XomeTox evaluating multi-omic integration for assessing rodent thyroid toxicity” has been set to investigate the utility of multi-omic data integration in toxicology and develop best practices aiming at regulatory applications^17^. In an application for drug safety, the recent study by Selevsek et al. used time-resolved proteome, transcriptome and methylome measurement in iPSC-derived human 3D cardiac microtissues to successfully elucidate adverse mechanisms of anthracycline cardiotoxicity^18^. In addition, the use of quantitative phosphoproteomics has been recently explored to compare distinct signaling events induced by four stressors with different modes of action^19^. Indeed, the idea of using phosphoproteomic alongside transcriptomic measurements to investigate cellular responses to different stressors has been proposed previously^20^. In particular, it has been hypothesized that the direct regulation of the activity of existing stress proteins through post-translational control could provide a faster adaptive cellular response to chemical exposure to ensure that the cell stays in a homeostatic state^20, 21^.

In this work we studied cellular responses to caffeine and coumarin by measuring and integrating multi-omic data generated from transcriptomics, proteomics and phosphoproteomics experiments. The two chemicals have been chosen due to their differences in the mechanism of action and because they have been the subject of two recent next generation risk assessment case studies^22, 23^. Caffeine is a compound whose mechanism of action is thought to be mediated via the following molecular initiating and key events: the antagonism of adenosine receptors, the inhibition of phosphodiesterase, the release of calcium from intracellular stores, and antagonism of benzodiazepine receptors^24–26^. One of the most sensitive mechanisms appears to be the reversible binding and inhibition of the adenosine A2A receptor^27^. In contrast, coumarin is a compound with a low bioactivity and its mechanism of action is not well-defined ^28, 29^. For both compounds, we generated proteomics, phosphoproteomics and transcriptomics concentration-response data respectively at two time points using a control and six treated HepG2 cell samples. The responses detected have been used to interrogate different levels of molecular activities in a dynamic fashion and capture an unprecedented level of details on the cellular responses elicited by increasing concentrations of the two chemicals.

## MATERIALS AND METHODS

### Chemicals and Reagents

Caffeine was purchased from TAUTO (Shanghai, China). Coumarin, the antibody against phospho-histone H2A.X (Ser139) (mouse), Trypsin-EDTA solution, and phosphatase inhibitor cocktails 2 and 3 were obtained from Sigma-Aldrich (St. Louis, MO, USA). The secondary fluorescent antibody Alexa Fluor 488 (goat anti-mouse) and ProLong™ Gold Antifade Mountant with DAPI were from ThermoFisher Scientific (Waltham, Massachusetts, USA). Dulbecco’s modified eagle medium (DMEM), fetal bovine serum (FBS), and penicillin-streptomycin were obtained from Gibco (Grand Island, NY, USA). Phosphate buffered saline (PBS) was acquired from HY Clone (Logan, Utah, USA). CellTiter-Glo® Luminescent cell viability assay kit was obtained from Promega (Madison, WI, USA). TRIzol reagent was obtained from Invitrogen (Carlsbad, CA, USA). SmartScribe RTase kit was acquired from Clontech (Mountain View, CA, USA). Phusion® High-Fidelity DNA Polymerase was from NEB (Beverly, MA, USA). EDTA-free protease inhibitor was purchased from Roche (Basel, Basel-Stadt, Switzerland).

### Cell Culture and Chemical Treatment

HepG2 cells (ATCC, Manassas, Virginia, USA) were cultured in DMEM medium supplemented with 10% FBS, 100 U/mL penicillin and 100 U/mL streptomycin at 37L in a humidified atmosphere of cell incubator containing 5% CO_2_. Cells used for all experiments were grown to approximately 80% confluence. Caffeine was dissolved in DMEM (100 mM stock solutions) and DMSO (1000 mM stock solutions), respectively. Further dilutions were made with serum-free DMEM. At the time of caffeine exposure, after the FBS-DMEM culture medium was removed, HepG2 cells were washed with PBS once and were finally treated in the fresh serum-free DMEM culture medium containing different concentrations of caffeine (0-1000 μM) for indicated time (10 min, 6 h, 24 h). The treatment with coumarin was performed using a dose range 0-100 μM for 10 min, 6 h and 24 h, as described previously^30^.

### Cytotoxicity Assays

Cytotoxicity assays included the cell viability assay and immunofluorescence for detecting DNA damage. The cell viability assay was performed according to the manufacturer’s instructions. Briefly, 4 × 10^3^ HepG2 cells were seeded per well into a 96-well culture plate for 12 h before treatment. Then, cells were treated in triplicates with 6 concentrations of caffeine (0.01, 0.1, 1, 10, 100, 1000 μM) and coumarin (0.001, 0.01, 0.1, 1, 10, 100 μM) which were serially diluted with serum-free DMEM for different time points (10 min, 6 h, 24 h). Following chemical treatment, the cells were further incubated at 37 L for 4 h after adding 20 μL MTS solution to each well. Finally, the absorbance was detected at 490 nm in the microplate reader. Data were normalized to time-matched control cultures which were considered as 100% cell survival.

The assay of immunofluorescence for detecting DNA damage was performed as described before^29^. Briefly, HepG2 cells were grown on microscope cover glass in 12-well culture plates with a seeding density of 8 × 10^4^ cells/well for 18 h. Afterwards, the cells were treated with 6 concentrations of caffeine and coumarin, which were the same as for the cell viability assay, for 6 h and 24 h. Then, the cells were fixed with 4% paraformaldehyde and permeabilized with 0.1% Triton X-100 in PBS at room temperature for 15 min and 5 min, respectively. After blocking in FBS for 1 h, the cells were incubated with the antibody against phospho-histone H2A.X (Ser139) (1:200 diluted in PBS) at 4 L overnight and subsequently incubated with the secondary fluorescent antibody Alexa Fluor 488 for 1 h at room temperature. Finally, the cells were measured with ProLong™ Gold Antifade Mountant with DAPI. After the mountant dried, the slides were examined at ×200 magnification with a confocal laser scanning microscope (ZEISS LSM 880, Carl Zeiss, Oberkochen, Baden-Württemberg, Germany). Each treatment had 3 biological replicates and the images were randomly captured.

### Transcriptomics, Proteomics and Phosphoproteomics Experiments

For transcriptomics, 5×10^7^ HepG2 cells were treated with a solvent control or 6 concentrations of caffeine for 6 h and 24 h before recovering. The experiments were performed in triplicate. Total RNAs were purified by Trizol reagent, quantified by a NanoPhotometer and converted to cDNA using SmartScribe RTase kit according to manufacturer’s protocol. The cleaned reads were then aligned to the genome reference of Ensembl GRCh38 with Hisat2 using strand specific parameters. For proteomics and phosphoproteomics, the same HepG2 cells were recovered immediately after 10 min and 24 h of caffeine treatment. Proteins were then digested in-gel with acetylated trypsin^31, 32^ and LysC at 37 °C for 14 h. Subsequently, peptides were enriched with the Ti^4+^-immobilized metal ion affinity chromatography (IMAC) method, as previously described^33, 34^. The peptides of total cell lysate and enriched phoshopeptides were separate into 3 fractions and then detected by Q-Exactive HF mass spectrometer (Thermo Fisher Scientific, San Jose, CA, USA) and LUMOS mass spectrometer (Thermo Fisher Scientific, San Jose, CA, USA) with a data dependent acquisition (DDA) mode. The raw data of mass spectrometry of proteomics and phosphoproteomics was parsed by MaxQuant software (version 1.6.0.1) using human database from UniProt (version 201506). For more details refer to the Supporting Information.

### Data Pre-filtering and Differential Expression Analysis

For RNAseq data, a typical pre-filtering criterion of median counts over all samples < 5 was carried out. The normalization and analysis of differentially expressed genes (DEGs) were conducted by using DESeq2^35^ which is widely used for differential expression analysis for experiments with fewer than 12 replicates^36^. For proteomics and phosphoproteomics, reverse and potential contaminant proteins and phosphosites with a localization probability < 0.75 were deleted. Proteins and phosphosites with intensities detected in at least 4 out of 21 samples were kept for further analysis. The data were then log_2_-transformed and median-normalized using Perseus software^37^. Subsequently, the missing values of the data were imputed by following the normal distribution implemented in Perseus software (see Supporting Information for more details).

For both caffeine and coumarin, DEGs, DEPs and DEPSs were identified by using a pre-filtering procedure with Benjamin-Hochberg adjusted p-value < 0.05 with different fold-change (FC) thresholds. The following FC thresholds were chosen: 1.5 for transcriptomics, 2 for proteomics, and 3 for phosphoproteomics. The choice of FC for each omic technique was informed by performing Gaussian fitting and deriving a 95% confidence value. For more information, refer to Figure S1 and Supporting Information Fold Change Thresholds.

### Dose Response Based on Bayesian BMD and Pathway Enrichment Analysis

For the concentration-response analysis of caffeine and coumarin, responsive genes, proteins, and phophosites were analyzed as continuous data by the Bayesian BMD method (BBMD)^38^ with the pre-processing algorithm of one-way ANOVA using a screening criterion of p-value < 0.05 and the FC thresholds as for the DE analysis. Data were then fitted to 7 different models: Exponential 2, Exponential 3, Exponential 4, Exponential 5, Hill, Power and Linear. The Benchmark Response (BMR) factor of 10% above/below the control level based on the average model was used to determine the POD at the gene/protein/phosphoprotein level (POD_T_/POD_P_/POD_Ph_). For further analyses, we focused on the features with reliable curve fits and meaningful BMD values, hence, the BMD analysis output was further filtered based on model average with BMD <= top experimental concentration and the ratio of BMDU/ BMDL < 40.

At the pathway level, the responsive pathways were selected based on the Gene Ontology (GO)^39^ and Reactome pathway^40^ databases containing at least 3 input genes with a p value < 0.05. The BMDL mean value of each pathway for transcriptomics and proteomics was calculated based on the average of BMDL values of the genes or proteins enriched in the particular pathway. For phosphoproteomics, the BMDL mean value of each enriched pathway was calculated using the phosphosite with the lowest BMDL value when there were multiple phosphosites showing response in one protein. Global POD values for each time point and each omic experiment were determined at pathway and gene level based on the previously published approaches^41^: (1) significantly enriched pathway with the lowest BMDL mean value: this approach revealed the pathway that was perturbed at the lowest dose. (2) BMDL mean value of 20 enriched pathways with the lowest p values: this method shows the pathways which may be most associated with chemical perturbation. (3) BMDL mean value of 20 pathways with lowest BMDL: this may reveal the most significantly responsive pathways after chemical treatment. (4) Mean of all pathway BMDLs. (5) Mean BMDL of features between 25th and 75th percentile: this may show the central measure of response to chemicals stressors.

### Multi-Omics Factor Analysis (MOFA)

In order to identify coordinated transcriptional and protein expression responses, MOFA analysis was performed using the transcriptome and proteome data at 24 h^42, 43^. Intuitively, MOFA can be viewed as a factor analysis applied to multi-omic data obtained on paired samples. The algorithm implemented as an R package is freely available on GitHub (https://github.com/bioFAM/MOFA2) and was used here to obtain MOFA factors and associated weights. These (latent) factors can be thought of as principal components and represent the driving sources of variation across the omic datasets. The factor weights provide a score for how strongly each feature relates to each factor.

### Kinase Prediction Analysis

Responsive phosphosites after 10 min exposure to caffeine and coumarin were used for kinase prediction analysis which was conducted using the iGPS software^44, 45^ in which the kinase-specific sequence motifs in substrates and protein-protein interactions (PPIs) between protein kinases and substrates were used to predict the upstream kinases. The threshold criterion was set as ‘high’ and the interaction filter was set as ‘Experimental/String’. BMD-filtered DEPSs were used as input for the analysis.

### Transcription Factor (TF) Analysis

Transcription factor enrichment analysis was performed with ChIP-X Enrichment Analysis 3 (ChEA3) ^46^. As input, we used BMD-filtered responsive genes from the 6 h transcriptomics experiments as these are likely more proximal to the molecular initiate events (MIEs) than the 24 h response. The output included top 10 TFs with the number of overlapping genes > 3, which were ranked using Integrated Scaled Rank method.

### Data Access

The RNA-seq data for caffeine has been uploaded to the Gene Expression Omnibus website (https://www.ncbi.nlm.nih.gov/geo/) with the accession number of GSE200441 and GSE200700. The mass spectrometry proteomics and phosphoproteomics data for caffeine have been uploaded to the ProteomeXchange Consortium website (http://proteomecentral.proteomexchange.org) via the iProX partner repository^47^ with the dataset identifier PXD030943, PXD030946, PXD031031, PXD031005. The transcriptomics, proteomics and phosphoproteomics data for coumarin can be found in our previous study^30^.

## RESULTS

### Experimental design

In order to identify dynamically altered transcripts, proteins and phosphosites upon caffeine and coumarin exposure, we compared the treatment time profiles with control profiles derived from time-matched DMSO-treated HepG2 cells. The choice of time points was informed by the response time characteristics of the individual omic technology. The 10 min time point was chosen to probe the start of an adaptation process detected in the phosphoproteome^17^. The 6 h time point in the transcriptomic experiment was previously found to be relevant to the direct MIE mechanism resulting from the chemical exposure^48, 49^. Lastly, both transcriptomics and proteomics were carried out for a longer exposure duration of 24 h to maintain consistency with our previous studies^5, 22, 23^.

### Caffeine and coumarin did not induce any significant changes in cytotoxicity up to 100 µM

For each time point, each omic experiment was carried out using 6 concentrations and the vehicle control using 3 biological replicates. In order to maintain consistency with the previous risk assessment studies of coumarin and caffeine^22, 23^, concentrations for caffeine treatment were preselected, ranging between 0.01-1000 µM at a 10-fold increment and coumarin ranging between 0.001-100 µM also at a 10-fold increment. The choice of concentrations was further confirmed here by measuring changes in cell count and DNA damage by using cell imaging with fluorescence analysis and immunocytochemistry of mouse mAb to phospho-Histone H2A.X Ser139 (Figure 1). No significant caffeine-induced cytotoxicity was observed up to 100 µM (Figure 1a-c). At 1000 µM, a drop in cell count of 20% was reported upon caffeine exposure for 6 h and 24 h, but not after 10 min. This drop in cell viability at 1000 μM was corroborated in the DNA damage experiment where HepG2 cells were exposed to caffeine at various concentrations for 6 h and 24 h (Figure 1g). Similarly, coumarin did not induce any significant changes in cytotoxicity up to a maximum tested concentration of 100 µM. A minor drop in cell count of 5% was observed at the exposure level of 100 µM for 24 h (Figure 1d-f). No DNA damage was observed at any of the tested concentrations of coumarin (Figure 1h).

**Figure 1.**
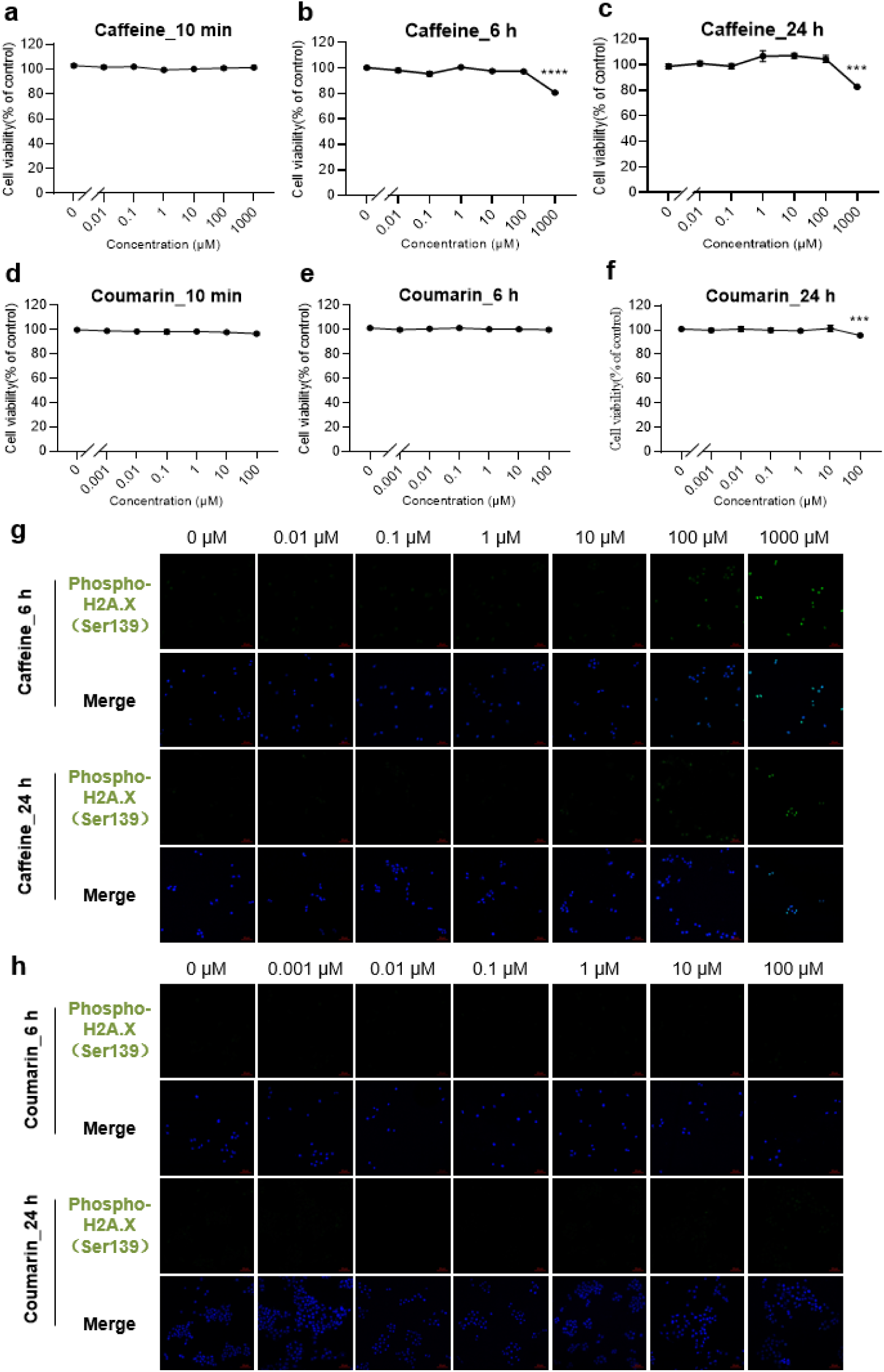
Cell viability and DNA damage upon treatment of HepG2 cells with caffeine and coumarin. Cell viability detected when cells were treated with caffeine and coumarin for 10 min (a & d), 6 h (b & e), and 24 h (c & f), respectively. Each measurement was done in 3 biological replicas. (g) DNA damage after the treatment of varied concentration of caffeine for 6 h and 24 h, as indicated. (h) DNA damage after the treatment of varied concentration of coumarin for 6 h and 24 h, as indicated.

### Quality Control of Transcriptomics, Proteomics and Phosphoproteomics Data

The quality control showed high quality of the data with the number of detected transcripts, proteins and phosphosites summarized in Table S1. The number of detected proteins (over 8,000) and phosphosites (over 47,000) were at the upper end of those previously reported^50, 51^ highlighting the improvements and breadth in coverage now possible for robust toxico-proteomic analysis. The pairwise Pearson correlation coefficient between the samples was relatively high. For more details on these and other quality control parameters, refer to Supporting Information. For the downstream analyses, we included only proteins and phosphorylation sites that were quantified in at least 4 out of 21 samples of a time point and for which the UniProt accession could be mapped to an Ensembl ID. As a result, 16,965 (6 h) and 16,729 (24 h) transcripts, 6,987 (10 min) and 6,988 (24 h) protein groups, as well as 27,680 (10 min) and 23,739 (24 h) phosphosites were used in the subsequent analyses for the caffeine treated samples (Table S1). For the coumarin treated samples, this resulted in 16,080 (for both 6 h and 24 h) transcripts, 6,230 (10 min) and 6,001 (24 h) protein groups, and 16,541 (10 min) and 14,157 (24 h) phosphosites (Table S1).

For a comprehensive statistical analysis of the dose response data, we used the Bayesian BMD (BBMD) software^38^. To limit the amount of data for subsequent model fitting, the data were first pre-filtered (see Materials and Methods Data Pre-filtering and Differential Expression Analysis) and hence restricted to features that showed some indication of a dose response trend. Thus, the number/fraction of features passing pre-filtering gave us a first rough estimate of how many transcripts/phosphosites/proteins were affected by the treatment with caffeine or coumarin in a dose-dependent manner, the so-called DEGs, DEPs and DEPSs. Using identified DEGs, DEPs and DEPSs, we then performed BMD modeling and functional classification analysis to determine the dose response behavior of the cells and infer the mechanism of action of caffeine and coumarin in HepG2 cells.

### HepG2 cells responded to caffeine exposure first by altering the phosphorylation status of existing proteins, then by changing expression levels of a number of genes which was followed by changes in protein abundance of a relatively small number of proteins

For all three omic experiments, the number of DEGs, DEPs and DEPSs increased with the increase in concentration of caffeine, as expected (Figure 2g). For the transcriptome (Figure 2a), the number of DEGs at lower concentrations of caffeine (0.01-10 µM) was on average higher at 6 h than at 24 h, suggestive that early responses had subsequently returned to baseline, whereas at higher doses (100-1000 µM) this effect was reversed. For the proteome, the number of detected DEPs was significantly larger at 24 h than at 10 min (Figure 2b). This behavior was expected given that 10 min represents a short timeframe for protein translation and synthesis, and thus abundance changes, to occur^17^. For the phosphoproteome, where a fast response is expected, early effects at 10 min were indeed stronger than those detected after 24 h and produced a higher number of DEPSs across all doses (Figure 2c). Lastly, when we compared the number of identified DEGs, DEPs and DEPSs at each assay concentration, it was evident that the number of DEPs did not follow the number of DEGs and only a very small fraction of identified proteins showed any kind of dose response after 24 h (Figure 2g).

**Figure 2.**
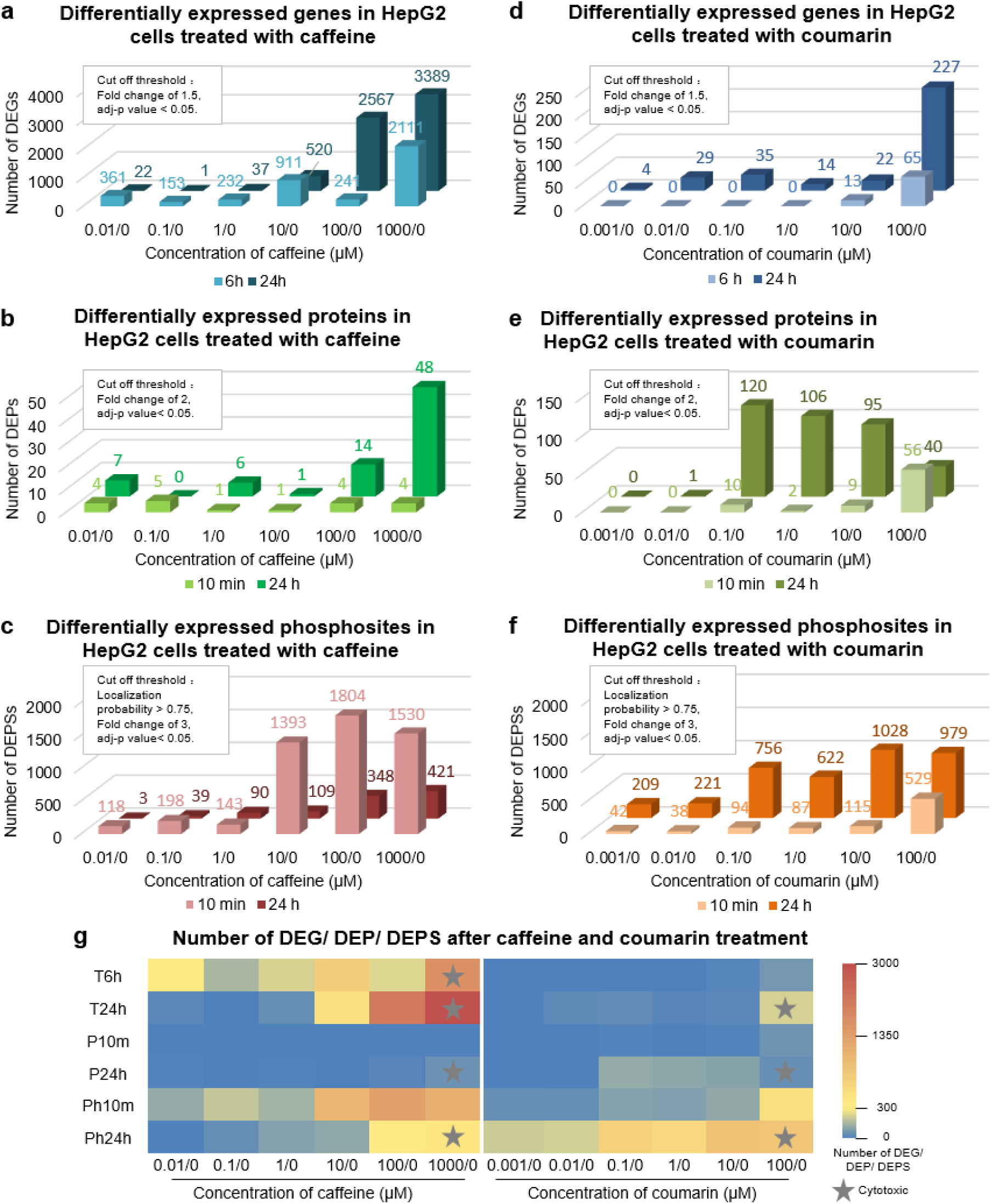
Number of detected DEGs, DEPs, DEPS upon caffeine and coumarin treatment. Number of DEGs (a), DEPs (b) and DEPSs (c) induced by caffeine treatment that passed the pre-filtering procedure as a function of time (rows) and concentrations (columns). Number of DEG (d), DEPs (e) and DEPSs (f) induced by coumarin treatment that passed the pre-filtering procedure as a function of time (rows) and concentrations (columns). (g) Heat map for the number of DEGs, DEPs and DEPSs. The gray star indicates the concentration at which the drop in cell viability was detected.

We then determined genes/proteins/phosphosites that showed a dose response to the caffeine exposure (see Materials and Methods Dose Response Based on Bayesian BMD and Pathway Enrichment Analysis). We first binned BMD values in appropriate concentration ranges between 0-1000 µM and looked at the number of responsive genes/proteins/phosphosites in each range across the three omic experiments (Figure 3a). From Figure 3a, it was evident that BMD_Ph_ values spanned a wide range of doses between 0-1000 µM. On the other hand, all BMD_T_ values laid above 1 µM and all BMD_P_ values were above 10 µM. When the total number of proteins that passed quality control was considered (6,987 for 10min and 6,988 for 24 h), it was striking that only a very small fraction of proteins had BMD_P_ values below 1000 µM; 55 and 45 proteins at 10 min and 24 h, respectively.

**Figure 3.**
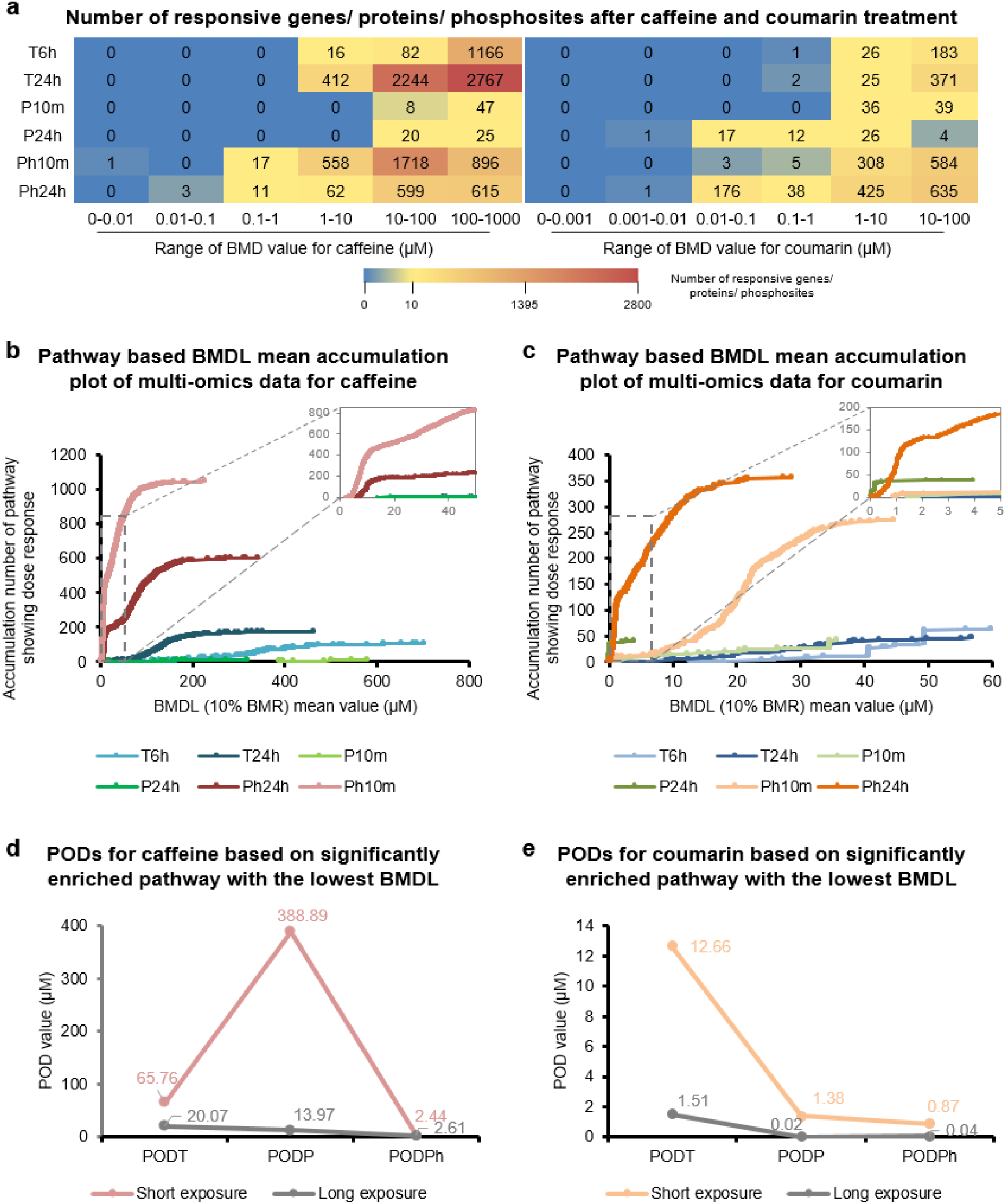
Number of responsive features based on BMD analysis. (a) Heat map with the number of BMD-responsive features binned according to the BMD value. BMDL mean accumulation plot for the multi-omics data for caffeine (b) and coumarin (c). Each data point represents the total number of GO terms and Reactome pathways that met the significance criteria (BMR = 10%, at least 3 input genes with p < 0.05 and a ratio of BMDU/BMDL < 40). Comparison of global POD values for caffeine (d) and coumarin (e). Short exposure for transcriptomics, proteomics and phosphoproteomics denoted 6 h, 10 min and 10 min, respectively. Long exposure denoted 24 h.

We then used BMDL_T_, BMDL_P_ and BMDL_Ph_ values to derive accumulation scores on the level of pathways and global POD_T_, POD_P_ and POD_Ph_ (Materials and Methods Dose Response Based on Bayesian BMD and Pathway Enrichment Analysis). Pathway accumulation scores, depicted in Figure 3b, represent a cumulative number of enriched pathways at a given concentration of caffeine derived from the gene/protein/phosphosite level BMDL_T_, BMDL_P_ and BMDL_Ph_. From Figure 3b, it is evident that the number of accumulated pathways coming from perturbations in the phosphoproteome at 10 min and 24 h was the highest (Figure 3b). This was followed by the accumulation number of enriched pathways coming from the transcriptional response at 24 h and 6 h. Of note, the accumulation number of enriched pathways coming from the proteome response was by far the smallest. Lastly, we derived global POD_T_, POD_P_ and POD_Ph_ using several published approaches that have been shown to correlate closely to standard pathological studies (Figure 3d, Figure S8)^41^. We found that the changes in the phosphoproteome happened at the lowest dose of caffeine with POD_Ph_ in the range 2.44-35.12 µM for the 10 min exposure and 2.61-71.32 µM for the 24 h exposure. This was followed by POD_T_ that spanned the range from 65.76 - 341.31 µM and 20.07 – 143.82 µM for the 6 h and 24 h exposure, respectively. POD_P_ was the highest and spanned the range from 388.89 - 485.53 µM and 13.97 – 160.93 µM for the 10 min and 24 h exposure data, respectively. Of note is that the total number of enriched pathways coming from the proteomics data was below the threshold value of 20 that is required by the global POD_P_ methods.

In summary, the presented results based on differential expression, BMD and POD analysis show that HepG2 cells adapt to the caffeine exposure by altering the phosphorylation status of existing proteins starting with the lowest dose of caffeine. This was followed by the gene expression that occurs at higher concentrations of caffeine (above 20 µM) and lastly a very small number of proteins was eventually altered at very high assay concentrations.

### HepG2 cells responded less strongly to the coumarin than caffeine exposure

The number of DEGs detected at 6 h was consistently lower than that at 24 h across all assay concentrations (Figure 2d). Of note, there were no DEGs detected at 6 h for the dose range between 0.01-1 µM. For the proteome, the number of detected DEPs was larger at 24 h than at 10 min (Figure 2e). This behavior was expected, as concluded for the caffeine data, given that 10 min represents a short timeframe for protein translation and synthesis to occur^17^. For the phosphoproteome, effects at both 10 min and 24 h were detected across the whole dose range (Figure 2f). Unlike for the caffeine dataset, the effects at 24 h were stronger than those detected at 10 min and produced a larger number of DEPSs across all doses (Figure 2f). When the number of identified DEGs, DEPs and DEPSs were compared at each assay concentration (Figure 2g), we found that the number of DEPs at 24 h, concentration range 0.1–10 µM was higher than the respective numbers of DEG at both 6 h and 24 h. It is uncertain whether this is a real effect or a consequence of a higher level of noise in the proteomics dataset at 24 h. Also, it is of note that the relative number of DEPSs at 10 min and DEGs at 6 h was much lower than the respective numbers found in the caffeine data. Because these time points are more proximal to the MIE direct effect, the reported numbers indicate a lower bioactivity of coumarin in HepG2s in relation to caffeine.

We then binned the BMD values in appropriate concentration ranges between 0-1000 µM and looked at the number of responsive genes/proteins/phosphosites in each bin across the three omic experiments (Figure 3a). Overall, it is evident that the numbers of coumarin BMDs across the concentration ranges were lower compared to the respective caffeine BMDs numbers (Figure 3a). This could be another indication of a generally lower bioactivity of coumarin in HepG2s than caffeine. Interestingly, the number of BMD_p_s found at 24 h for the range 10-100 µM was lower than the respective numbers found for bins 0.01–0.1, 0.1–1 and 1-10 µM. This may be a consequence of the adaptation process or a higher level of noise in the proteomics dataset for coumarin exposure at 24 h.

Consistent with the differential expression and BMD analysis, the overall number of enriched pathways depicted by the accumulation plot from the coumarin dataset was lower than that for the caffeine dataset (Figure 3c). This is another indication of a lower bioactivity of coumarin than caffeine. Again, the only exception was seen in the accumulation number of enriched pathways coming from the proteomic response at 24 h. Interestingly, when BMDL_T_s, BMDL_P_s and BMDL_Ph_s were used to derive the global POD_T_, POD_P_ and POD_Ph_ with the five methods described above^41^, it was consistently found that PODs coming from all coumarin omic experiments were lower than those coming from the caffeine experiments (Figure 3e, Figure S8). This is in contrast to the findings from the differential expression, BMD and pathway analysis. It is of note that the methods used for calculation of global PODs^41^ rely on either a single significantly enriched pathway or an average of a subset of pathways/genes and hence could be affected by the remaining level of noise in the coumarin dataset that had not been filtered out.

In summary, it was found that HepG2 cells adapt to the coumarin exposure by altering the phosphorylation status of existing proteins starting with the lowest dose used, similar to caffeine.

Based on the differential expression, BMD and pathway analysis we found that HepG2 cells responded less strongly to the coumarin than caffeine exposure. Inconsistencies in the global POD values were likely a consequence of the higher level of noise present in the coumarin dataset.

### A more comprehensive understanding of pathway responses to chemical exposure could be obtained by integrating multi-omics data

#### Caffeine

To examine a potential mechanism of action of caffeine, we next examined compound-responsive features and their associated over-/under-represented GO [6] terms and Reactome pathways [7]. First, we looked at the overlap between the dose response features to examine whether the affected features are (di)similar across the omic datasets (Figure 4a-b, Figure S9a-b, Table S2-S5). As shown in Figure 4a-b and Figure S9a-b, we found a very few overlapping features, GO terms and Reactome pathways, similar to conclusions reported previously^18^. Most of the inter-omic overlap was between the DEPSs and DEGs, and in particular between the phosphoproteome at 10 min and transcriptome at 6 h (223 features and 101 GO terms/Reactome pathways). This was expected given that the phosphorylation signal can directly or indirectly activate/deactivate the transcription of relevant genes^21^. However, when we looked at the overlap of the BBMD analysis output, we observed hardly any inter-omic overlap; the overlap was intra-omic mainly between the phosphoproteome at 10 min and 24 h and between the transcriptomes at 6 h and 24 h.

**Figure 4.**
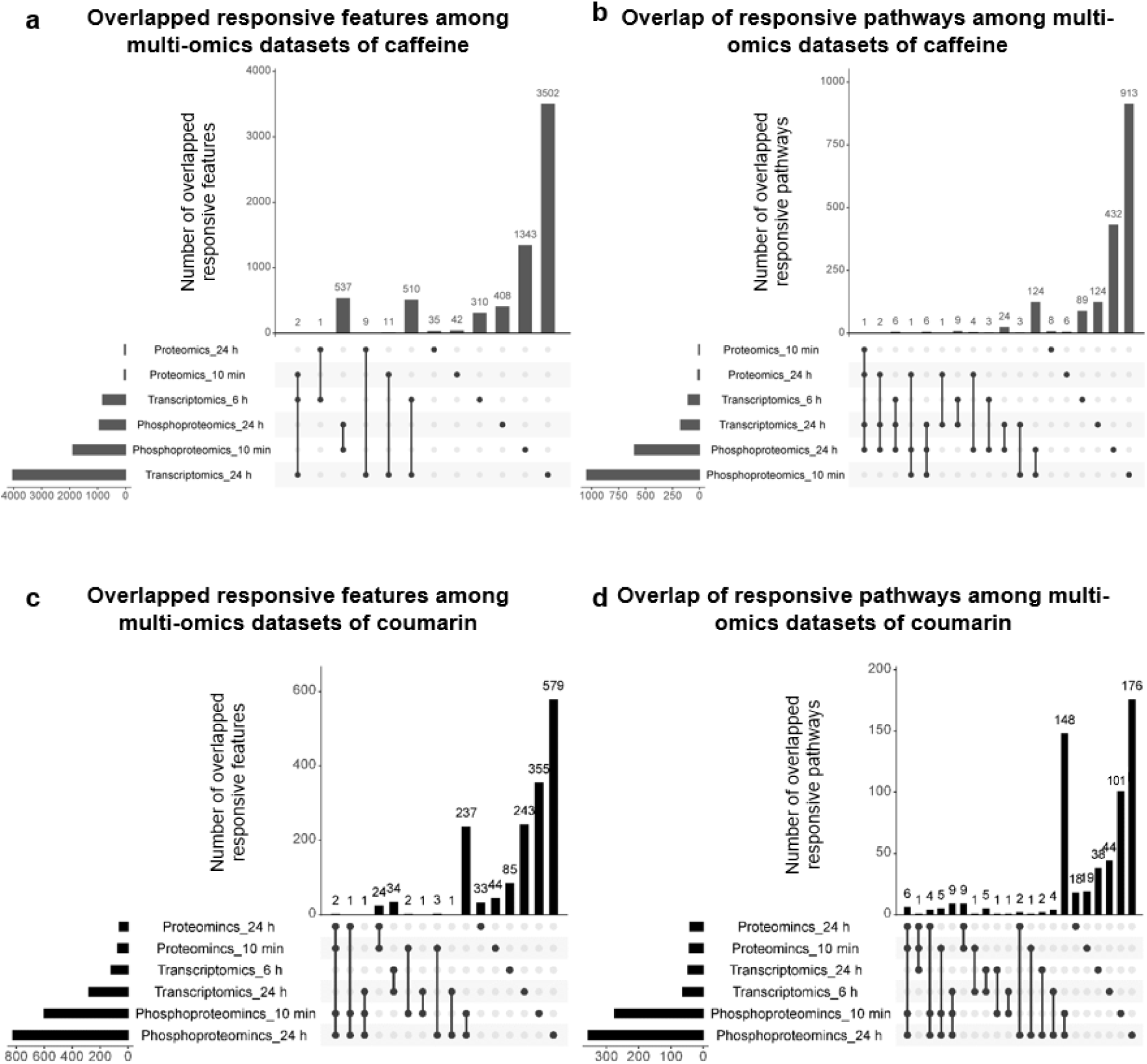
Overlap of BMD-filtered dose responsive features between the different omic datasets. Overlap of individual features for the caffeine (a) and coumarin (c) dataset. Overlap of GO terms and Reactome pathways for the caffeine (b) and coumarin (d) dataset. The upset plot was generated using an online platform (http://www.bioinformatics.com.cn).

**Figure 5.**
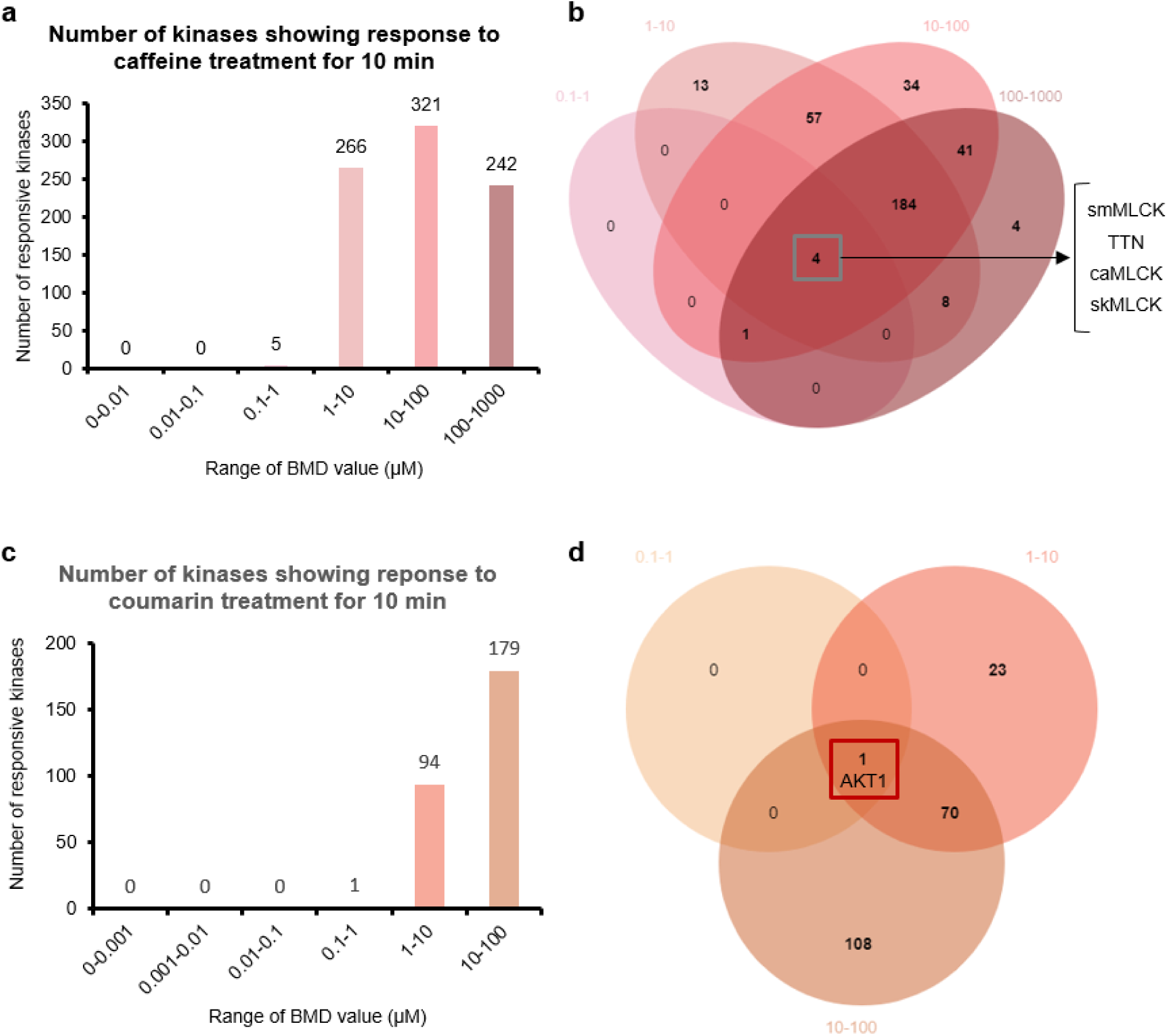
Number of upstream kinases identified from the responsive phosphosites after 10 min exposure at different BMD values. The number of kinases identified from the caffeine (a) and coumarin dataset (c). Overlap of the kinases identified between treatment groups from the caffeine (b) and coumarin (d) dataset.

We then looked at top 20 significantly enriched GO terms and Reactome pathways coming from the DEGs and DEPs (Figure S10-S12, Table S6). Our focus was on the terms and pathways at the relatively low level of hierarchy as those were particularly useful when considering the putative mechanism of action. From the transcriptomics data at 6 h, the general transcription factor activity had the lowest p-value across GO terms and Reactome pathways. In particular, the *genes expression regulated by PPARalpha pathway* was found as the most enriched Reactome pathway (lowest p-value). Interestingly, unsaturated fatty acid biosynthetic/metabolic process related to *PPARalpha pathway* was found to have the lowest p-value in the proteomics dataset at 24 h but was not among top enriched pathways in the transcriptomics data at 24 h. This indicated that the proteins that regulate fatty acid metabolism were indeed synthesized in response to the caffeine exposure and were found enriched after 24 h of exposure. Again, no overlap between GO terms and Reactome pathways was found when the analysis was performed on filtered BMD features (Table S7). Findings that caffeine induces fatty acid turnover and lipid oxidation have been reported in the past^52–54^. Also, it is of particular note that the same mechanism of action in HepG2 cells has been reported before when triglyceride and cholesterol levels were measured upon treating the cells with caffeine^55^.

We then looked whether multi-omic integration can further inform on the common biology present in our omic datasets. For that we used MOFA analysis that is able to integrate paired samples, herein the transcriptomics and proteomics data at 24 h for both caffeine and coumarin (see Materials and Methods Multi-Omics Factor Analysis)^42, 43^. Briefly, MOFA infers a low-dimensional representation of the multi-dimensional data in terms of a small number of latent factors that capture the global sources of variability. As shown in Figure S13a-d, most of the signal from the proteomics and transcriptomics datasets was encoded by Factor 1 where cholesterol biosynthesis (proteomics 24 h) and general process of DNA replications (transcriptomics 24 h) were found as the most overrepresented pathways. This finding was in agreement with the functional analysis carried out on the level of each omic experiment.

Lastly, we examined whether the phosphoproteome and transcriptome can inform on the upstream signaling events as a result of caffeine exposure (see Materials and Methods Kinase Prediction and TF Analysis). To this extent, we identified upstream protein kinases based on BMD filtered DEPSs and transcription factors based on BMD filtered DEGs. We were mainly interested in upstream events proximal to the MIEs and hence we only considered the phosphoproteome at 10 min and transcriptome at 6 h. As shown in Figure 5a-b and Table S8, we were able to identify 834 kinases with BMDs in range from 0.1 to 1000 µM. Among the detected kinases, titin (TTN) and 3 isoforms of the myosin light chain (MLC) kinase, namely, skeletal, smooth muscle and cardiac were detected across all assay doses – with all four kinases being implicated in the regulation of contractions in skeletal, smooth and cardiac muscle. Indeed, it has previously been reported that caffeine interacts with MLC kinase either directly by causing relaxation of smooth muscle^56, 57^ or indirectly by causing vasodilation^58^. It has also been reported that healthy and TTN-related cardiomyocytes differ in their responses to caffeine^59^.

In addition, we were able to identify top 10 ranked TFs with BMDs in range 10–100 µM and 100–1000 µM (see Table S9 for details). Among the TFs identified for the BMD range 10-100 µM, there were two TFs belonging to the FOX family (FOXD4L4, FOXD4L1) that have previously been implicated in developmental processes^60^ and four TFs belonging to the zinc finger family (ZNF526, ZNF556, ZNF776, ZNF793) that have previously been implicated in a wide range of cellular processes^61^. Among the TFs identified for the BMD range 10-1000 µM, there were three nuclear receptors CAR, PXR and RORC that are well-known regulators of the metabolism of a variety of endogenous and xenobiotic compounds. It is of note, that we did not identify AhR among the top TFs, despite the fact that AhR regulates CYP1A2 which is responsible for the metabolism of caffeine^62^. This could be because the base expression level of CYP1A2 is below the level of detection in HepG2 cells^63^ which was also confirmed in our transcriptomic dataset of both untreated and treated samples.

#### Coumarin

Similarly, the coumarin dataset showed a very few overlapping dose responsive features, GO terms and Reactome pathways (Figure 4c-d, Figure S9c-d, Table S10-S13). Unlike the caffeine dataset, there was barely any overlap between different omic experiments for both dose response features and the filtered BMD analysis output (Figure 4c, Figure S8c).

We then looked at top 20 significantly enriched GO terms and Reactome pathways coming from DEGs and DEPs (Figure S14-16, Table S14). Again, our focus was on the terms and pathways at the relatively low level of hierarchy as those are particularly useful when considering the putative mechanism of action. From the transcriptomics data at 6 h, we found no significantly enriched GO terms or Reactome pathways at the low level of hierarchy. On the other hand, the transcriptomics data at 24 h showed signs of general cell stress and was enriched for pathways such as *Cellular response to stress*, *Cellular response to stimuli* and *Unfolded protein response*. Furthermore, we found *Response of EIF2AK4 (GCN2) to amino acid deficiency* among top pathways in both transcriptomics and proteomics dataset at 24 h. This pathway is also an indicator of cellular stress and acts by reducing translation of most mRNAs and activation of genes involved in responding to amino acid deficiency. Again, no overlap between GO terms and Reactome pathways was found when the analysis was performed on filtered BMD features (Table S15).

We then looked whether multi-omic integration can further inform on the common biology present in the omic datasets using the MOFA analysis^42, 43^. As shown in Figure S13 e-h, Factor 2 encoded most of the signal coming from the proteomic dataset and Factor 1 from the transcriptomic dataset. In line with the previous findings, the majority of pathways coming from Factor 1 transcriptomics were related to general cellular stress. Among the enriched protein/gene sets encoded by both factors, we again found *Response of EIF2AK4 (GCN2) to amino acid deficiency* as the only pathway in common.

Lastly, we examined whether the phosphoproteome and transcriptome can inform on the upstream signaling events as a result of coumarin exposure. As shown in Figure 5c-d and Table S16, we were able to identify 274 kinases as upstream regulators of coumarin induced changes in the phosphoproteome at 10 min. Among them, the AKT1 kinase was the only kinase found across all doses. AKT1 is implicated in a wide range of processes including metabolism, proliferation, cell survival, growth and angiogenesis. To our knowledge, there have been no reports to indicate activation/deactivation of AKT1 as a response to coumarin treatment of HepG2 cells.

In addition, we were able to identify top 10 ranked TFs with BMDs in range 10–100 µM (see Table S17 for details). Among the TFs identified, there were two TWIST-related proteins (TWIST1, TWIST2) that have previously been implicated in cell differentiation and developmental processes^64^ and four TFs belonging to the zinc finger family (ZNF425, ZNF575, ZNF596, ZNF805) that are well-known regulators of a wide range of cellular processes^61^. It is of note that the main enzyme responsible for metabolism of coumarin CYP2A6^22^ is not expressed in HepG2 cells^63^ which was also confirmed in our transcriptomic dataset of both untreated and treated samples. Therefore, it is of no surprise that TFs that have previously been implicated in the regulation of CYP2A6, such as HNF-4alpha, C/EBPalpha, C/EBPbeta, and Oct-1^65^, were not identified among the top ranked TFs.

## DISCUSSION

In summary, we cross-compared changes in the transcriptome, proteome and phosphoproteome in response to caffeine and coumarin treatment of HepG2s. Caffeine and coumarin have fundamentally distinct mechanisms of action and were used to understand commonalities and differences in the biomolecular response measured by the three omic technologies. We demonstrated that alterations in the phosphoproteome were detectable as early as after 10 min of exposure to both caffeine and coumarin. Such a fact occurred not only earlier in time, comparing to the proteome changes measured at 10 min, as expected, but also at much lower chemical concentrations in comparison to the proteome and transcriptome level. Indeed, the regulation at the phosphoproteome level was detected even at the lowest doses of exposure and reinforces the potential for posttranslational modification to act as a fast energy-efficient adaptive response with aim to keep the cell in homeostasis^20^. We demonstrated that the early 10 min phosphoproteomic response to caffeine was relatively larger to that to coumarin and resulted in 834 kinases identified as upstream catalysts of the detected (de)phosphorylation changes. Among the detected kinases, TTN and 3 isoforms of the MLCK kinase, namely, skeletal, smooth muscle and cardiac were detected across all doses – with both kinases being implicated in the regulation of muscle contraction. Indeed, it has previously been reported that caffeine either activates these kinases directly or indirectly^56–59^. On the other hand, we identified 274 kinases as upstream regulators of coumarin induced changes in the phosphoproteome at 10 min. Among them, the AKT1 kinase implicated in a wide range of processes including metabolism, proliferation, cell survival, growth and angiogenesis was the only kinase detected across all the doses; however, we could not find any previous study that confirms activation of AKT1 induced by coumarin.

We studied overlap in differentially expressed genes/proteins and pathways as a result of the chemical exposure. The observed changes in protein expression were found to a considerably lower extent than that seen for the transcript expression, which is consistent with the fact that protein abundance depends on localization and post-translational modification besides levels of mRNA^13–16^. For caffeine data, we identified metabolism of lipids and inhibition of *PPARalpha* as commonly enriched pathways in the transcriptomics response at 6 h and proteomics response at 24 h. Interestingly, findings that caffeine induces fatty acid turnover, lipid oxidation and hepatic lipid mechanism have been reported many times before^52–55^, but only further targeted testing can confirm that these phenomena are induced by the cellular mechanism identified here. Conversely, from the coumarin dataset we were not able to identify any common pathways in the transcriptomic and proteomic response. This could be a consequence of a very broad coumarin activity on the cell stress that has been hypothesized before^29^.

The presented analysis showed that integration of different omic layers to be used in toxicology is not straightforward and indeed poses many challenges. While our primary focus was on extracting commonalities between the three omic layers, the overlap across the datasets was fairly low. One of the challenges represented the choice of time points for different experiments. The choice of time points used in this study was based on previous knowledge: the 10 min time point was chosen to probe the start of an adaptation process detected in the phosphoproteome^17^; the 6 h time point in the transcriptomic experiment was previously found relevant to the direct MIE mechanism coming from the chemical exposure^48, 49^; the 24 h time point was chosen to probe paired transcriptomic and proteomic samples when using a longer exposure duration. However, we are aware that this is a complex issue and there is not only a time variance between different omic layers, e.g. proteome is downstream of the transcriptome, but also different biological processes occur on different time scale within a single omic layer, e.g. translation occurs on a different time scale from cell division^17^. In addition, since proteomics and phosphoproteomics have not been routinely used as NAM-based tools in hazard evaluation, the reproducibility of the data acquisition and consistency in the data analysis demand more consideration. Furthermore, the coverage of expressed proteins, phosphosites and transcripts needs to be addressed in order to ensure that relevant biology has been sufficiently covered. For example, the gene expression of the enzymes responsible for the metabolism of caffeine and coumarin was found below the level of detection in HepG2s, and that might have precluded the identification of the relevant transcriptional regulators of these enzymes. In addition, the HepG2 cells were found to express the well-known targets for caffeine such as adenosine receptor 1 (ADORA1) and adenosine receptor 2A (ADORA2A) below the level of detection. Therefore, any cellular response coming from the inhibition of the two receptors could not be detected in this study. Caffeine is also known to induce other effects, such as lipid metabolism which was detected here but the use of more relevant cell lines or a range of different cell lines might be necessary in future.

## CONCLUSIONS

HTTr is an omic technology that has already found application in characterizing the biological activity of chemicals in a non-animal risk assessment^5, 6, 41, 66^. Indeed, an increasing number of studies have demonstrated that HTTr can provide a broad biological coverage^23^, PoDs derived from HTTr are protective of human health^5, 67^ and can be used for decision making when combined with other NAMs^22, 23^(*in vitro* assays, PBK models). However, despite the protective nature of its PoDs, it is not completely clear if and how the very early biological perturbations detected by HTTr are linked to the functional adversity in the cell or tissue caused by the chemical exposure. For that reason, we have explored here whether other multi-omic experiments used alongside HTTr, such as proteomics and phosphoproteomics, can offer a more complete understanding of the perturbed toxicity pathways. We have shown that the methodology presented here is able to capture the complete chain of events from the first compound-induced changes at the phosphoproteome level to changes in gene expression induced by transcription factors and lastly to changes in protein abundance that further influence adversity on the cellular level. Phosphoproteomic changes were detected not only early in time but at very low exposure to both chemicals and hence are very relevant for studying proximal MIE effects induced by chemical exposure. On the other hand, changes in protein abundance were found much less frequently than the transcriptomic changes and can be used to further understand the role that the detected transcriptomic changes play in the pathway response to the chemical exposure. To our knowledge, this is the first in-depth study that measures cumulative effects on transcriptome, proteome and phosphoproteome induced by chemical exposure in a time and dose-dependent manner. Finally, as multi-omic experiments become more affordable, the application of such an approach to a larger number of chemicals, test conditions and cell lines will play an increasing role in building capability, understanding and ultimately confidence in bioactivity characterization in toxicity testing.

## AUTHOR INFORMATION

**Corresponding Author**

**Present Addresses**

### Author Contributions

P.X. and J.L. initiated the project and oversee all aspects of the project. F.X., K.L., S.L., S.Z.L. and Y.L. performed the experiment. Y.L. and P.X. performed all data analyses with the help of P.K., T.L., Z.Z., S.J., M.L., P.L.C. and A.W.. J.L., Q.Z., L.C., F.T. and X.G. assisted in promoting the project. Y.L., P.K. and P.X. wrote the manuscript with input from all the authors. All authors have given approval to the final version of the manuscript.

## ACKNOWLEDGMENT

We are indebted to Dr Shuangqing Peng, Dr Chao Liu and Dr Liang Zhao for support in the early stage of this project. We thank Dr Yangjun Zhang and Dr Guibin Wang for the help with reagents, peptide manual checking and LC-MS analysis of proteomics and phosphoproteomics samples. This study was funded by the Unilever 21th Century Toxicity Program (MA-2018−02170N), MOST (2017YFA0505100 and 2020YFE0202200), the National Natural Science Foundation of China (NSFC) (32141003, 31870824, 91839302, 31901037, 32070668 and 32071431), the Foundation of State Key Lab of Proteomics (SKLP-K201704, SKLP-K201901) and the CAMS Innovation Fund for Medical Sciences (2019-I2M-5-017), the Beijing-Tianjin-Hebei Basic Research Cooperation Project (J200001), the Mass Spectrometry Platform Open Project of National Center for Protein Sciences Beijing and Unilever-Emory Collaborative Project (MA-2015-02026).

## SUPPLEMANTARY FIGURES

**Figure S1.**
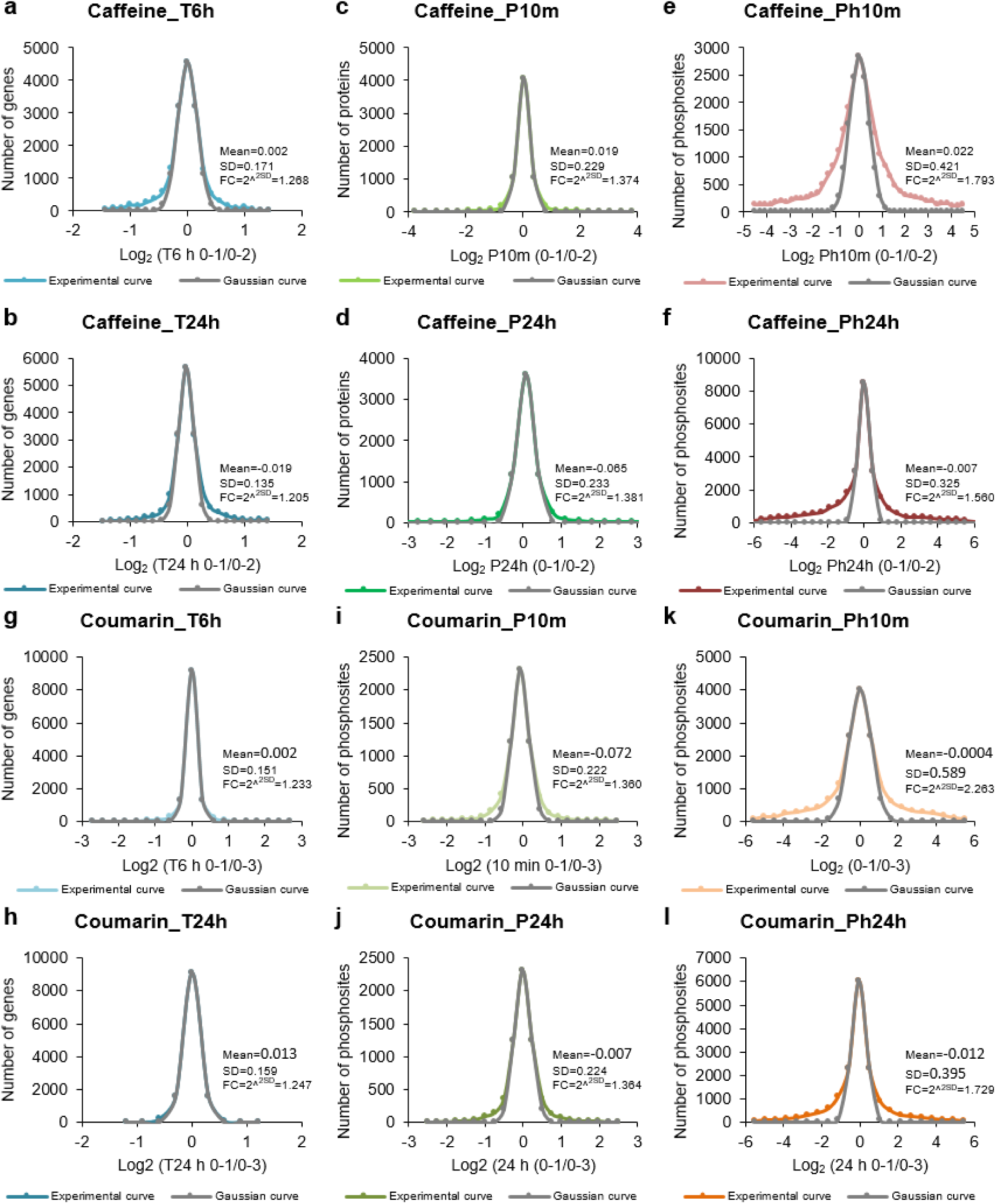
Distribution of the detected gene counts (transcriptomics) and intensities (proteomics, phosphoproteomics). The distributions were derived using 2 samples of 3 biological replicates in the vehicle control group of transcriptomics (a, b, g, h), proteomics (c, d, i, j), and phosphoproteomics data (e, f, k, l) of caffeine and coumarin and then compared to the Gaussian curve.

**Figure S2.**
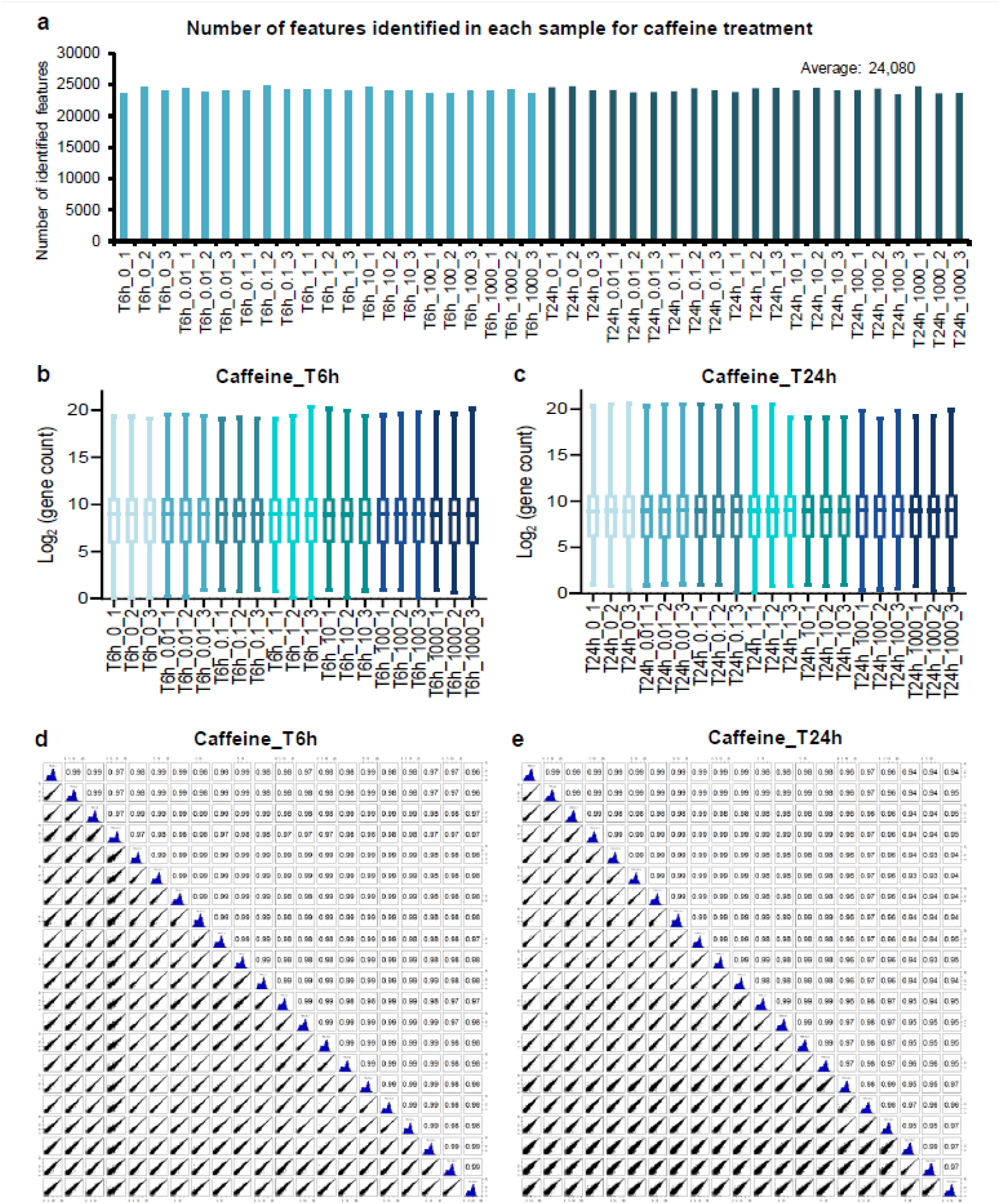
Quality control analysis for the caffeine transcriptomics dataset. (a) The number of identified features in each sample. Number of gene counts for each sample treated for 6 h (b) and 24 h (c). The Pearson correlation analysis between samples treated for 6 h (d) and 24 h (e).

**Figure S3.**
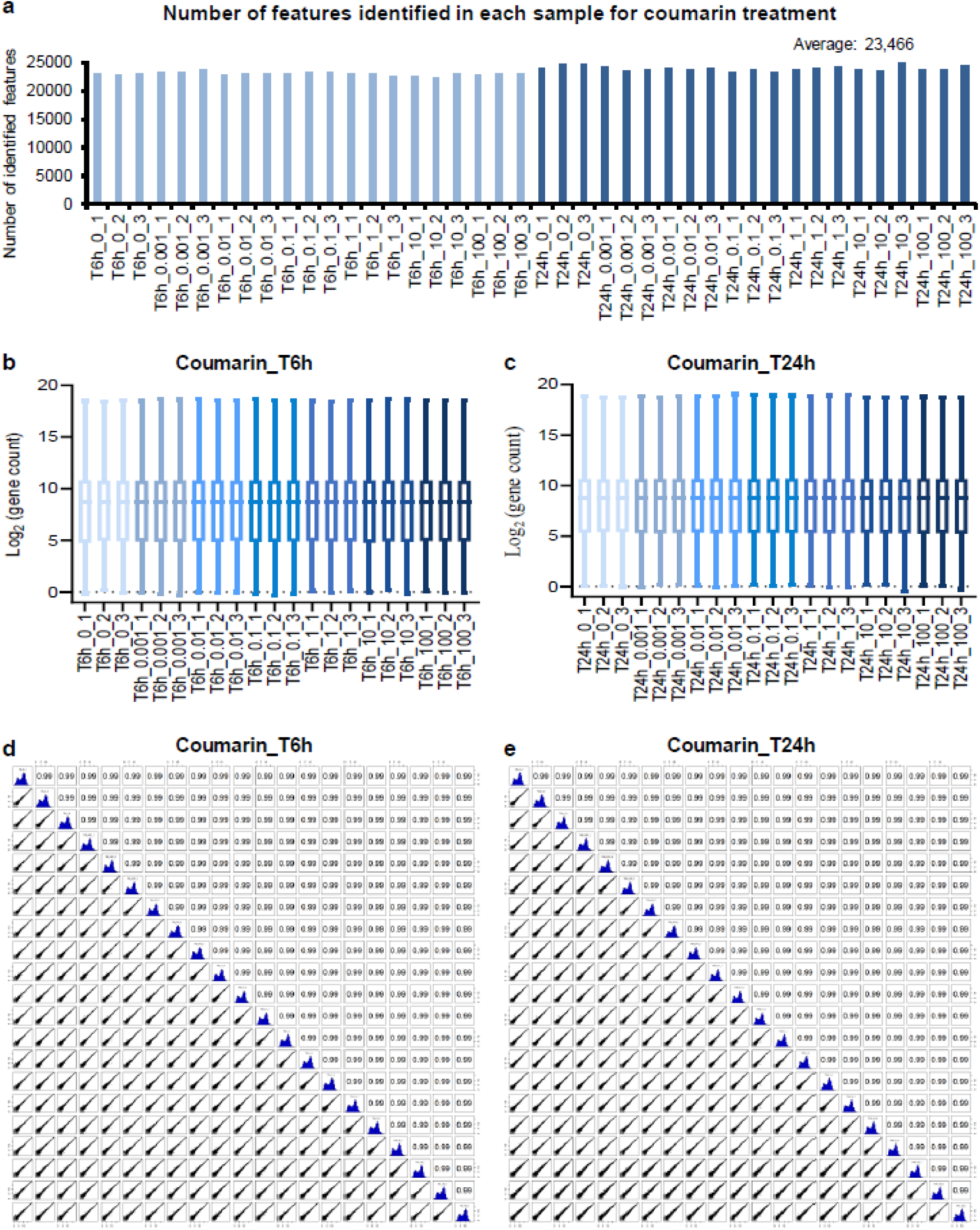
Quality control analysis for the coumarin transcriptomics dataset. (a) The number of identified features in each sample. Number of gene counts for each sample treated for 6 h (b) and 24 h (c). The Pearson correlation analysis between samples treated for 6 h (d) and 24 h (e).

**Figure S4.**
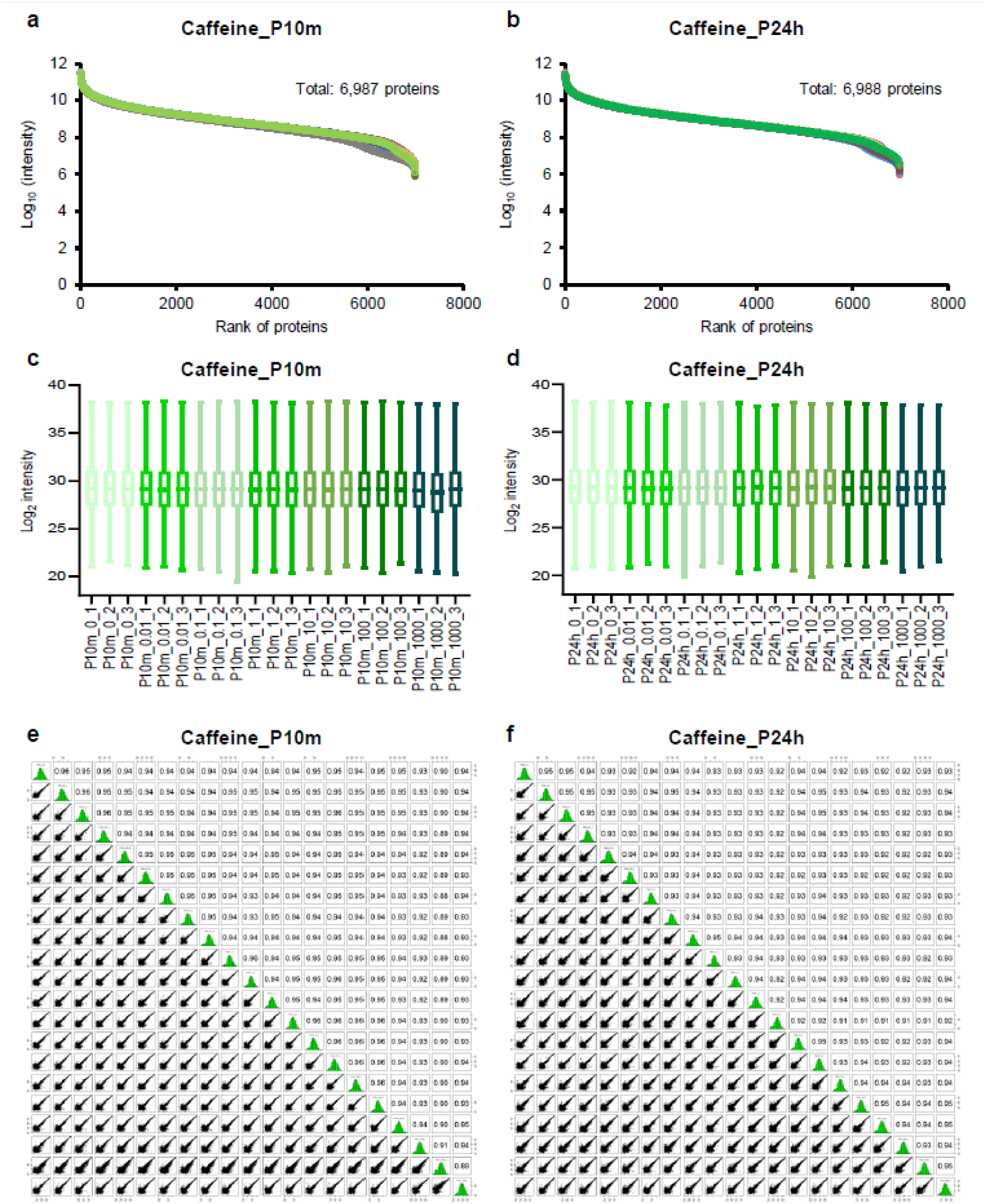
Quality control analysis for the caffeine proteomics dataset. Rank of identified proteins when the samples were treated for 6 h (a) and 24 h (b). Protein intensities for sample treated for 6 h (c) and 24 h (d). The Pearson correlation analysis between samples treated for 6 h (e) and 24 h (f).

**Figure S5.**
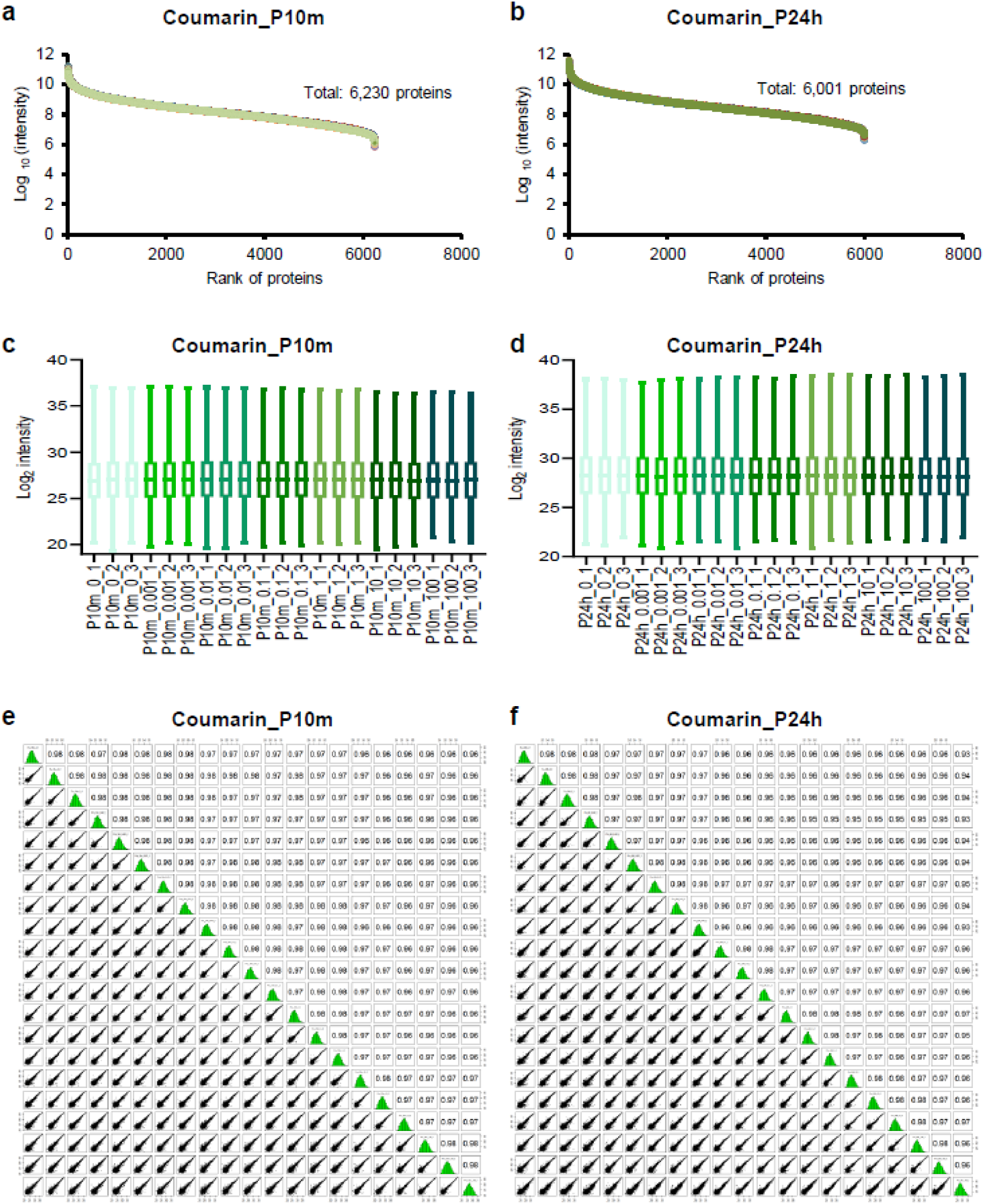
Quality control analysis for the coumarin proteomics dataset. Rank of identified proteins when the samples were treated for 6 h (a) and 24 h (b). Protein intensities for sample treated for 6 h (c) and 24 h (d). The Pearson correlation analysis between samples treated for 6 h (e) and 24 h (f).

**Figure S6.**
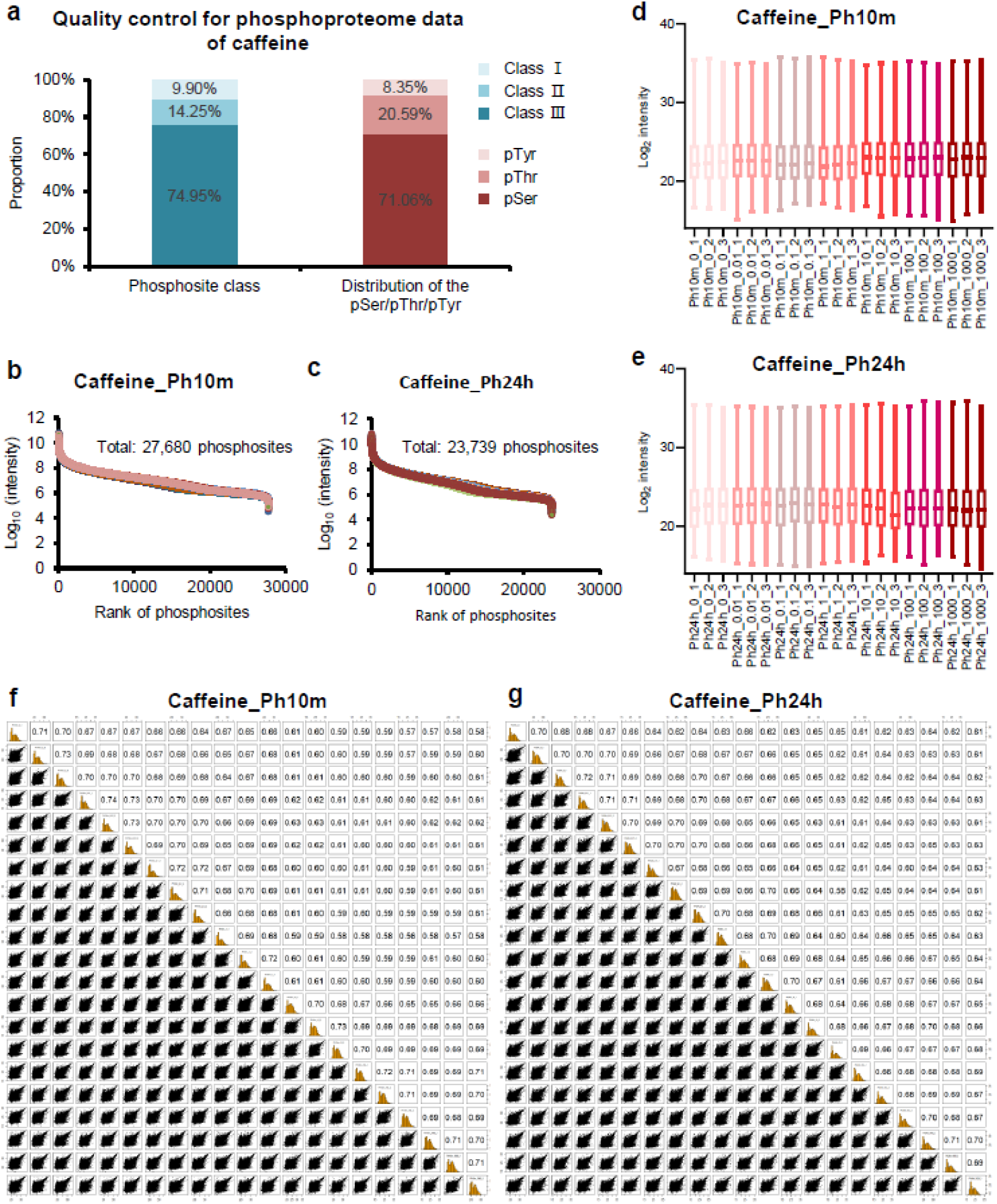
Quality control analysis for the caffeine phosphoproteomics dataset. (a) Distribution of the localization confidence for identified phosphosites (blue): Class ☐ (low confidence, probability of 0.25-0.5), Class ☐ (medium confidence, probability of 0.5 -0.75), Class ☐ (high confidence, probability >0.75). Distribution of the phosphorylated amino acids (red). Rank of identified phosphosites when the samples were treated for 6 h (b) and 24 h (c). Protein intensities for samples treated for 6 h (d) and 24 h (e). The Pearson correlation analysi between samples treated for 6 h (f) and 24 h (g).

**Figure S7.**
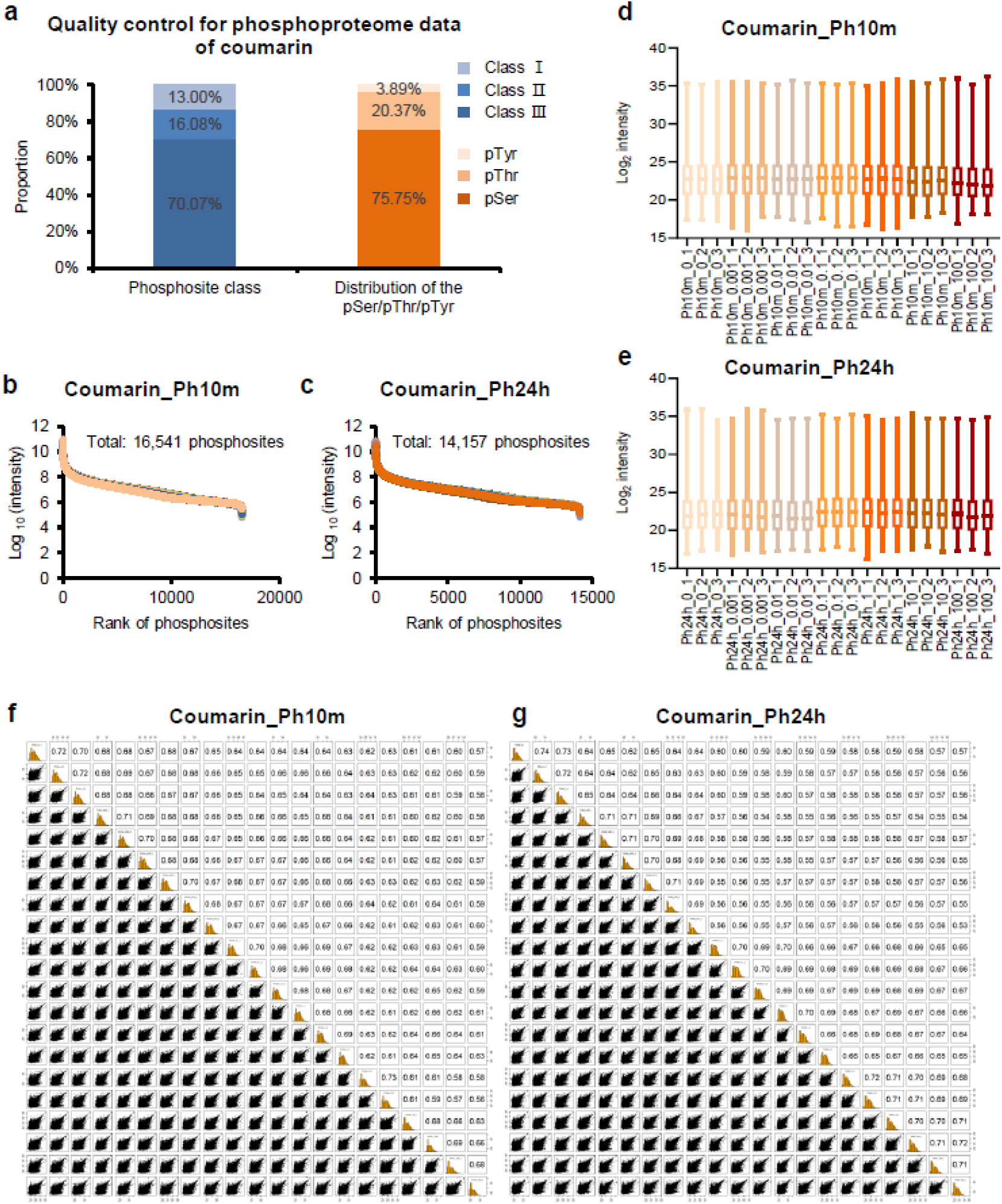
Quality control analysis for the coumarin phosphoproteomics dataset. (a) Distribution of the localization confidence for identified phosphosites (blue): Class L (low confidence, probability of 0.25-0.5), Class L (medium confidence, probability of 0.5 -0.75), Class L (high confidence, probability >0.75). Distribution of the phosphorylated amino acids (red). Rank of identified phosphosites when the samples were treated for 6 h (b) and 24 h (c). Protein intensities for samples treated for 6 h (d) and 24 h (e). The Pearson correlation analysis between samples treated for 6 h (f) and 24 h (g).

**Figure S8.**
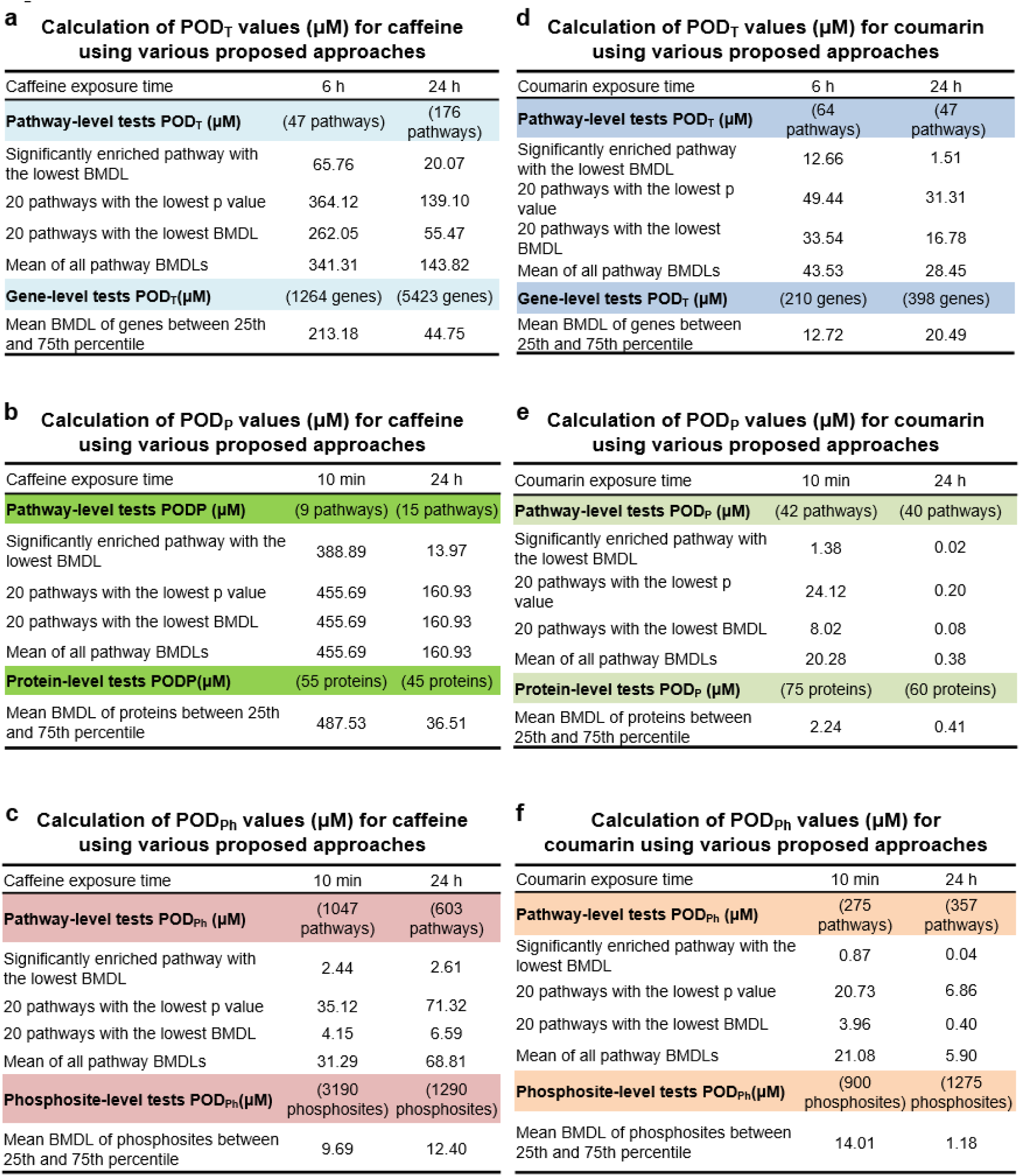
Comparison of POD_T_, POD_P_ and POD_Ph_ using the approaches detailed in Materials and Methods. POD_T_ (a), POD_P_ (b) and POD_Ph_ (c) values from the caffeine dataset. POD_T_ (d), POD_P_ (e) and POD_Ph_ (f) values from the coumarin dataset.

**Figure S9.**
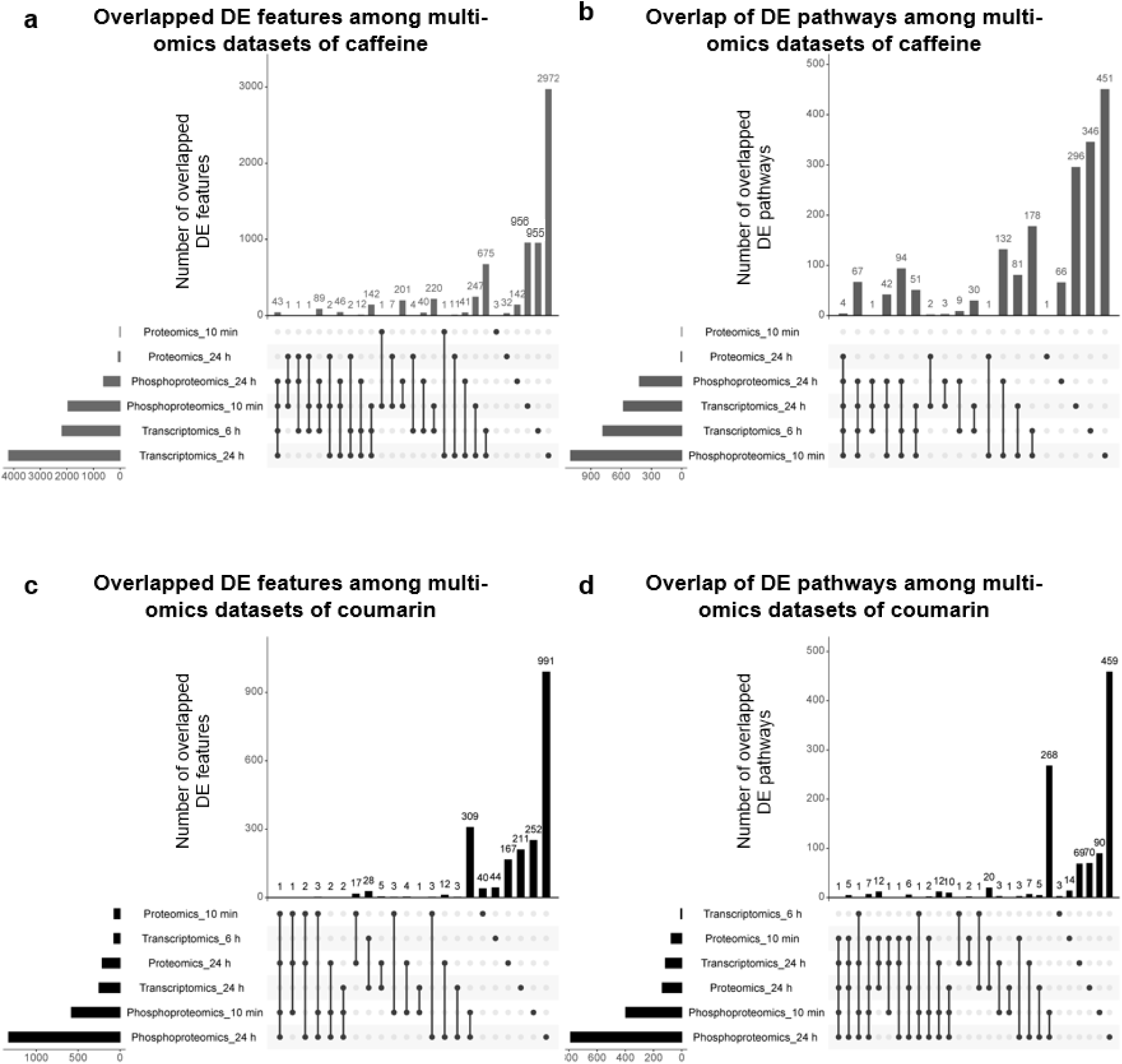
Overlap of differentially expressed features between the different omic datasets. Overlap of individual features for the caffeine (a) and coumarin (c) dataset. Overlap of GO term and Reactome pathways for the caffeine (b) and coumarin (d) dataset. The differentially expressed features were determined to be those with adj-p<0.05 with fold change of 1.5 for transcriptomics, 2 for proteomics, 3 for phosphoproteomics. The upset plot was generated using http://www.bioinformatics.com.cn, a free online platform for data analysis and visualization.

**Figure S10.**
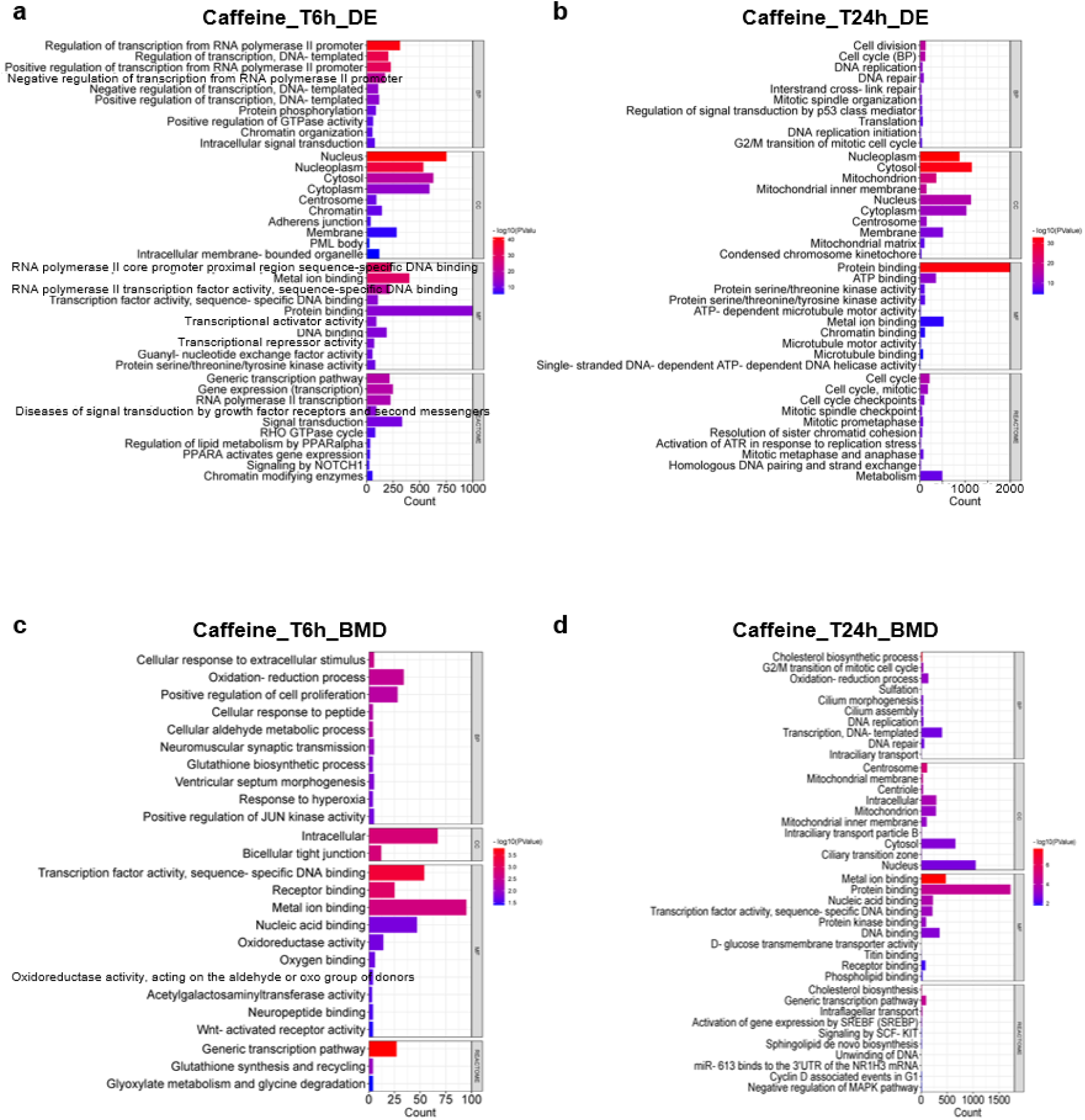
Functional analysis of the caffeine dataset based on GO terms and Reactome pathways using DEGs and BMD-filtered DEGs. Top 10 significantly enriched GO terms and Reactome pathways with the lowest p-value after the treatment for 6 h (a) and 24 h (b) based on the DEGs. Top 10 significantly enriched GO terms and Reactome pathways with the lowest p-value after the treatment for 6 h (c) and 24 h (d) based on the BMD-filtered DEGs. Each GO term and Reactome pathway was required to contain at least 3 genes with p < 0.05.

**Figure S11.**
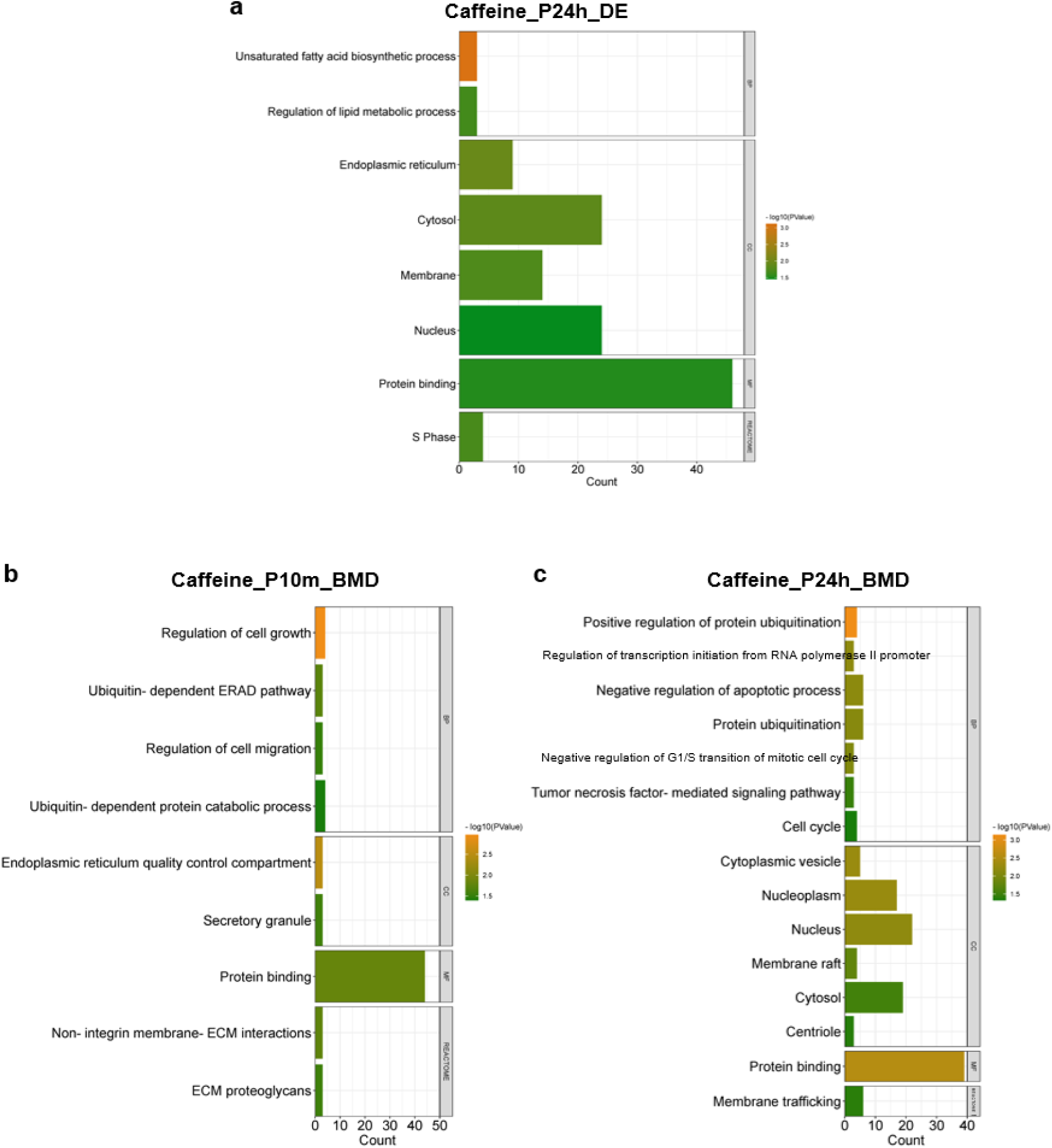
Functional analysis of the caffeine dataset based on GO terms and Reactome pathways using DEPs and BMD-filtered DEPs. Top 10 significantly enriched GO terms and Reactome pathways with the lowest p-value after the treatment for 24 h (a) based on the DEPs. There were no enriched pathways for the 10 min dataset. Top 10 significantly enriched GO term and Reactome pathways with the lowest p-value after the treatment for 10 min (c) and 24 h (d) based on the BMD-filtered DEPs. Each GO term and Reactome pathway was required to contain at least 3 genes with p < 0.05.

**Figure S12.**
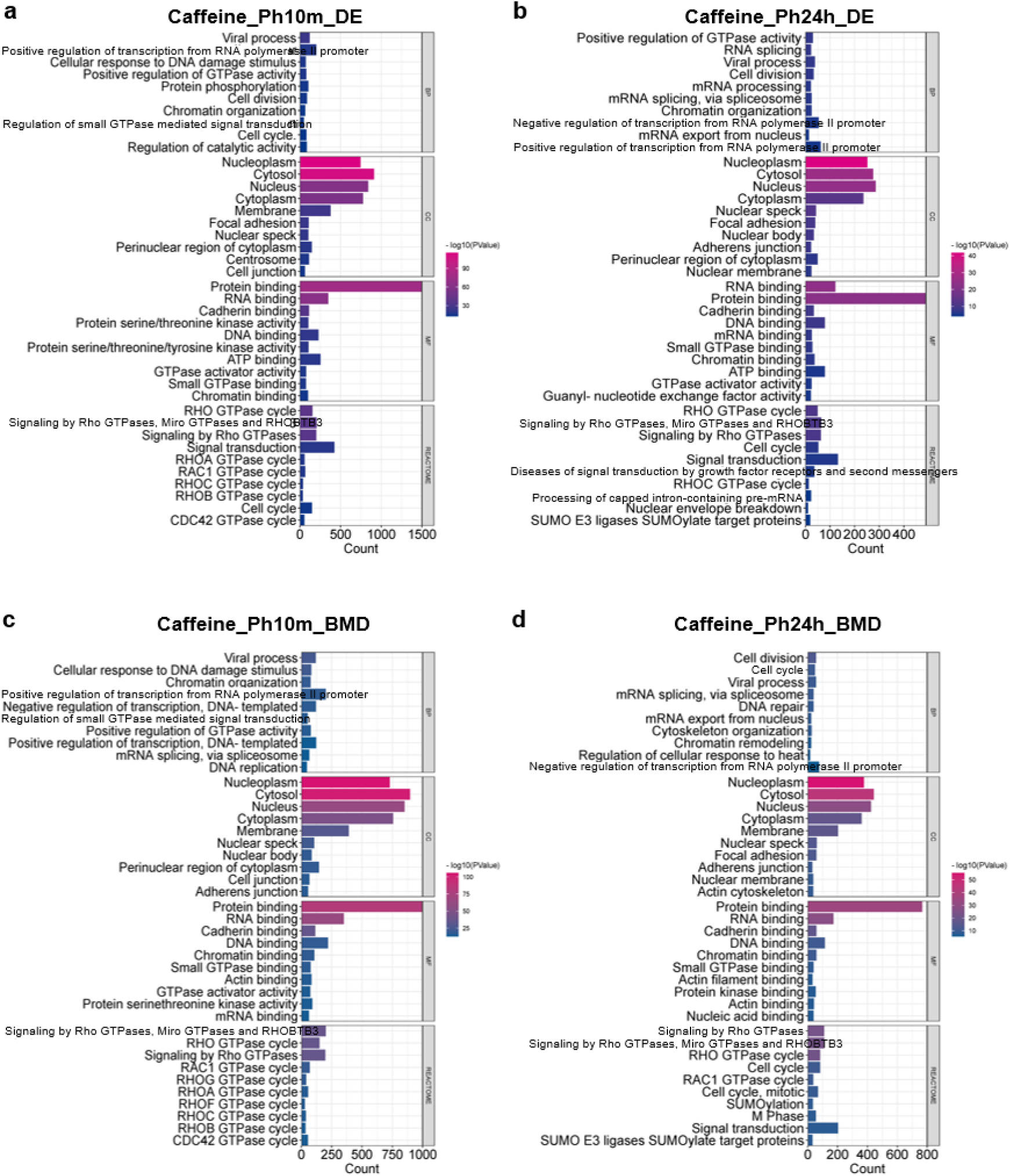
Functional analysis of the caffeine dataset based on GO terms and Reactome pathways using DEPSs and BMD-filtered DEPSs. Top 10 significantly enriched GO terms and Reactome pathways with the lowest p-value after the treatment for 6 h (a) and 24 h (b) based on the DEPSs. Top 10 significantly enriched GO terms and Reactome pathways with the lowest p-value after the treatment for 6 h (c) and 24 h (d) based on the BMD-filtered DEPSs. Each GO term and Reactome pathway was required to contain at least 3 genes with p < 0.05.

**Figure S13.**
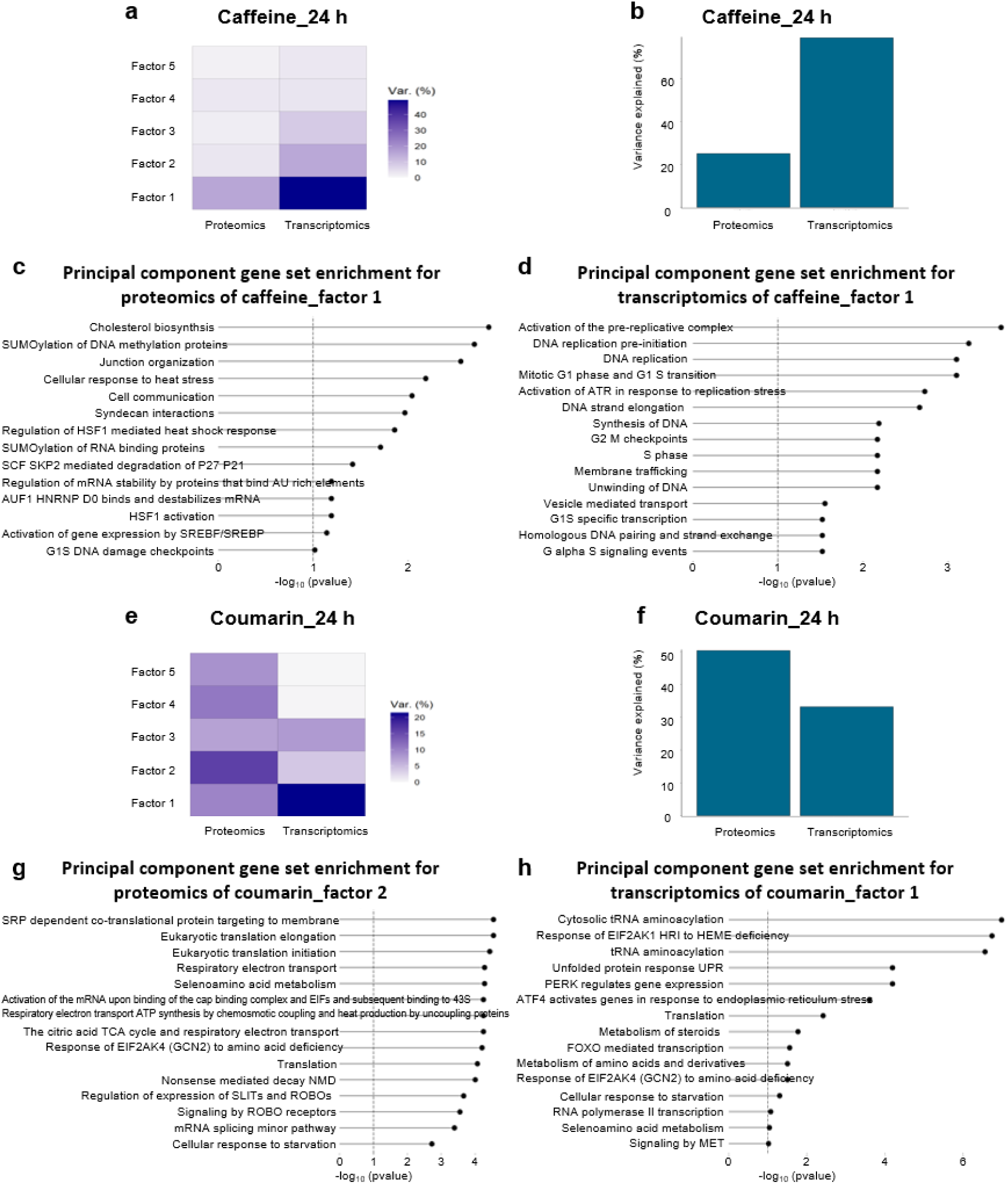
Multi-omic factor analysis (MOFA) using 24 h transcriptomics and proteomics dataset for caffeine and coumarin. (a-b) Variance explained by the MOFA model trained on the 24 h caffeine. Principal component gene set pathway enrichment for: factor 1 using the 24 h caffeine proteomics dataset (c) factor 1 using the 24 h caffeine transcriptomics dataset (d). (e-f) Variance explained by the MOFA model trained on the 24 h coumarin. Principal component gene set pathway enrichment for: factor 2 using the 24 h coumarin proteomics dataset (g) factor 1 using the 24 h coumarin transcriptomics dataset (h).

**Figure S14.**
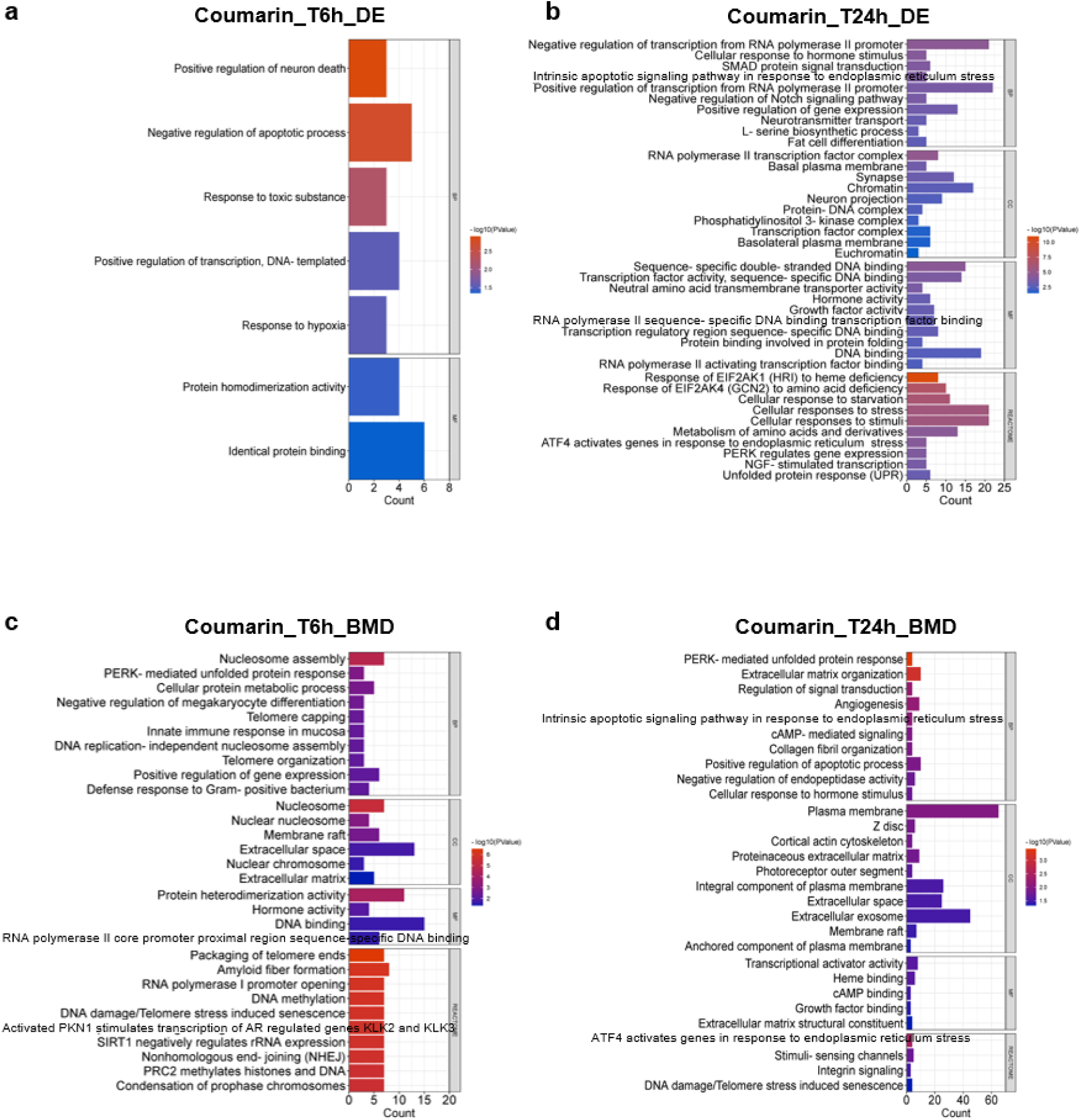
Functional analysis of the coumarin dataset based on GO terms and Reactome pathways using DEGs and BMD-filtered DEGs. Top 10 significantly enriched GO terms and Reactome pathways with the lowest p-value after the treatment for 6 h (a) and 24 h (b) based on the DEGs. Top 10 significantly enriched GO terms and Reactome pathways with the lowest p-value after the treatment for 6 h (c) and 24 h (d) based on the BMD-filtered DEGs. Each GO term and Reactome pathway was required to contain at least 3 genes with p < 0.05.

**Figure S15.**
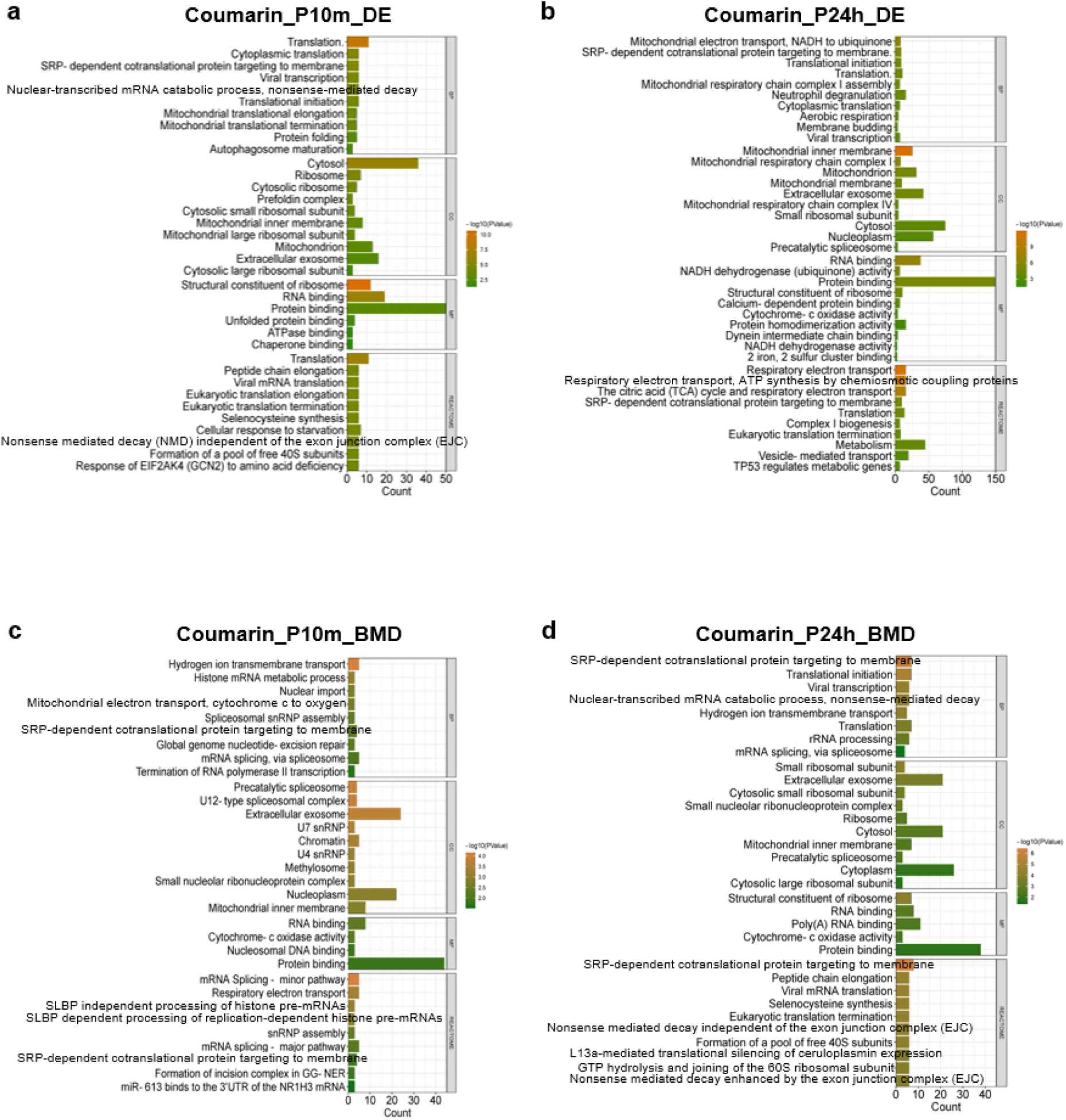
Functional analysis of the coumarin dataset based on GO terms and Reactome pathways using DEPs and BMD-filtered DEPs. Top 10 significantly enriched GO terms and Reactome pathways with the lowest p-value after the treatment for 6 h (a) and 24 h (b) based on the DEPs. Top 10 significantly enriched GO terms and Reactome pathways with the lowest p-value after the treatment for 6 h (c) and 24 h (d) based on the BMD-filtered DEPs. Each GO term and Reactome pathway was required to contain at least 3 genes with p < 0.05.

**Figure S16.**
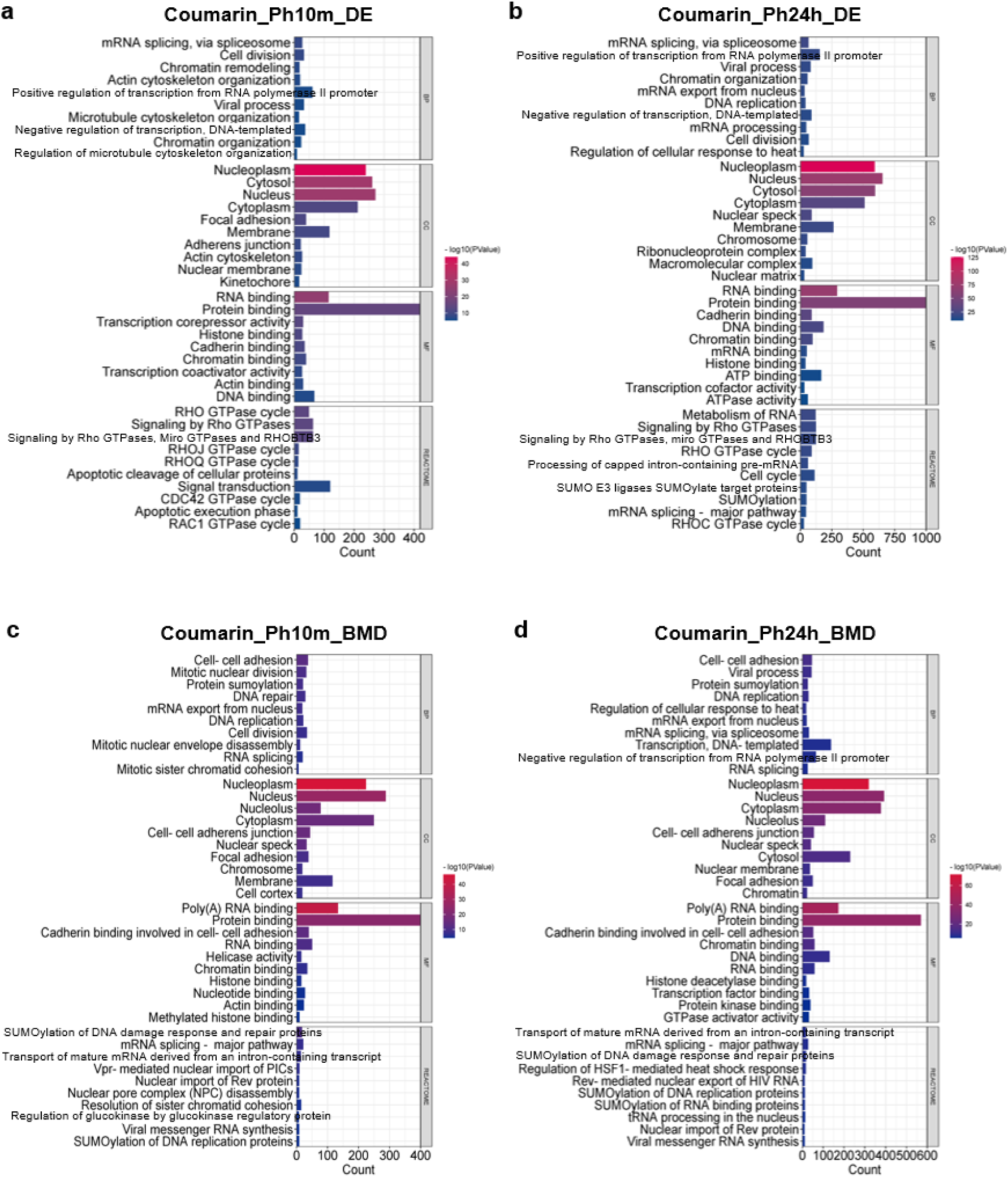
Functional analysis of the coumarin dataset based on GO terms and Reactome pathways using DEPSs and BMD-filtered DEPSs. Top 10 significantly enriched GO terms and Reactome pathways with the lowest p-value after the treatment for 6 h (a) and 24 h (b) based on the DEPSs. Top 10 significantly enriched GO terms and Reactome pathways with the lowest p-value after the treatment for 6 h (c) and 24 h (d) based on the BMD-filtered DEPSs. Each GO term and Reactome pathway was required to contain at least 3 genes with p < 0.05.

## SUPPLEMANTARY TABLES

**Table S1.** The number of detected transcripts, proteins and phosphosites.

**Table S2.** Overlap between differentially expressed features from the caffeine multi-omic dataset.

**Table S3.** Overlap between GO terms and Reactome pathways from the caffeine multi-omic dataset based on the differentially expressed features.

**Table S4.** Overlap between BMD-filtered dose responsive features from the caffeine multi-omic dataset.

**Table S5.** Overlap between GO terms and Reactome pathways from the caffeine multi-omic dataset based on the BMD-filtered dose responsive features.

**Table S6.** Top 20 significantly enriched GO terms and Reactome pathways from the caffeine multi-omic dataset based on the differentially expressed features.

**Table S7.** Top 20 significantly enriched GO terms and Reactome pathways from the caffeine multi-omic dataset based on the BMD-filtered dose responsive features.

**Table S8.** Upstream kinases identified from the responsive phosphosites after caffeine exposure for 10 min at different BMD values.

**Table S9.** Upstream TFs identified from responsive genes after caffeine exposure for 6 h at different BMD values.

**Table S10.** Overlap of differentially expressed features from the coumarin multi-omic dataset.

**Table S11.** Overlap of GO terms and Reactome pathways from the coumarin multi-omic dataset based on the differentially expressed features.

**Table S12.** Overlap of BMD-filtered dose responsive features from the coumarin multi-omic dataset.

**Table S13.** Overlap of GO terms and Reactome pathways from the coumarin multi-omic dataset based on the BMD-filtered dose responsive features.

**Table S14.** Top 20 significantly enriched GO terms and Reactome pathways from the coumarin multi-omic dataset based on the differentially expressed features.

**Table S15.** Top 20 significantly enriched GO terms and Reactome pathways from the coumarin multi-omic dataset based on the BMD-filtered dose responsive features.

**Table S16.** Upstream kinases the responsive phosphosites after coumarin exposure for 10 min at different BMD values.

**Table S17.** Upstream TFs identified from responsive genes after coumarin exposure for 6 h at different BMD values.

