## Supplemental materials for "Using NAMs to characterize chemical bioactivity at the transcriptomic, proteomic and phosphoproteomic levels"

This file of supporting information contains:

Pages: 8

Workflow of multi-omics experiments: page S2-S4

Quality control of multi-omics datasets: page S5-S7

Abbreviations: page S7-S8

References of supporting information: page S8

**WORKFLOW OF MULTI-OMICS EXPERIMENTS**

**Transcriptomics Experiment.** For transcriptomics, about 5×10^7^ HepG2 cells were treated in triplicates using 6 concentrations of caffeine and a control for 6 h and 24 h. After the culture medium with caffeine was removed, the cells were harvested by digestion of trypsin-EDTA solution and washed with ice-cold PBS twice and then centrifuged at 12,000 g for 15 min at 4 °C. RNAs were then purified by Trizol reagent. RNAs were quantified by a NanoPhotometer. The RNA samples were purified by poly (dT) beads, converted to cDNA using SmartScribe Rtase kit according to manufacturer’s protocol. Cleaned reads were aligned to the genome reference of Ensembl GRCh38 with Hisat2 using strand specific parameters. Raw RNAseq data for coumarin was obtained from our previous study^1^ where the identical experimental procedure was followed.

**Proteomics and Phosphoproteomics Experiment**

**Protein Extraction of Whole Cell Lysate.** For proteomics and phosphoproteomics, about 5×10^7^ HepG2 cells were treated in triplicates using 6 concentrations of caffeine and a control for 6 h and 24 h. As phosphoproteomics changes would be expected to occur shortly in seconds to minutes after exposure to chemicals^2, 3,^ we measured the samples after being treated for 10 min. In addition, samples were measured after being treated for 24 h and this time point was chosen as a representative of the steady state. After the culture medium containing caffeine was removed, HepG2 cells at approximately 80% confluency in 10 cm dish were harvested by digestion of trypsin-EDTA solution and washed with ice-cold PBS twice to get the cell lysis which was subsequently put in the freshly prepared lysis buffer (9 M Urea, 10 mM pH 8.0 Tris–HCl, 30 mM NaCl, 5 mM IAA, 5 mM Na_4_P_2_O_7_, 100 mM pH 8.0 Na_2_HPO_4_, 1 mM NaF, 1 mM Na_3_VO_4_, 1 mM sodium glycerophophate, 1% phosphatase inhibitor cocktail 2 and 3 and EDTA-free protease inhibitor). The cell lysis was placed on ice and sonicated by ultrasonic cell disruptor for 4 min (2-s on and 4-s off, amplitude 25%). After centrifuging (14,800 g) for 15 min at 4 °C to remove the cell debris, the supernatant was collected and stored at -80 °C for later use.

**Protein Digestion by Gel-aided Strategy.** Protein concentration was determined by a gel-assisted method as previously described^4^. For each sample, equal amount of 1 mg proteins was used for later procedures. Dithiothreitol (DTT) was added to cell lysis to a final concentration of 10 mM for 30 min at 45 °C. After that, the cysteine was alkylated by incubation with iodoacetamide (IAA) to a final concentration of 20 mM in the dark at room temperature for 30 min. Proteins were then digested in-gel with acetylated trypsin^5, 6^ at 37 °C for 14 h. After the digestion reaction was stopped, the tryptic peptides were collected by adding extracting buffer (5% formic acid (FA), 50% acetonitrile (ACN)), dried with a vacuum dryer and then stored at -80 °C for further analysis. In subsequent studies, 5% of the peptides were used for proteomics study and the remaining 95% for phosphoproteomics study.

**Phosphopeptide Enrichment by Ti^4+^-IMAC.** Peptides were enriched with the Ti^4+^-immobilized metal ion affinity chromatography (IMAC) method, as previously described^7, 8^. Briefly, the peptides were dissolved in 500 μL binding buffer (80% ACN, 6% TFA) and 500 μL loading buffer (10% 500 mM NH_4_HCO_3_, 5% ACN) followed by adding the Ti^4+^-IMAC beads with a ratio of peptides:beads = 1:25 (m/m). After vortexed severely for 30 min at room temperature, the mixtures of peptides and beads were centrifuged at 17,000 g for 6 min and the supernatant was removed. Then, the mixtures of peptides and beads were washed with 1.8 mL wash buffer 1 (50% ACN, 6% TFA in 200 mM NaCl) once for 30 min and 1.8 mL wash buffer 2 (30% ACN, 0.1% TFA in ddH_2_O) twice (30 min/time) to remove the non-specific adsorption of peptides. Finally, the phosphopeptides were eluted with 1 mL elution buffer (10% NH_3_·H_2_O) for 15 min, and then sonicated in ice water for another 15 min. The mixtures were centrifuged at 17,000 g for 6 min, and the supernatant was transferred into a new tube and lyophilized immediately.

**Fractionation of Peptides of Total Cell Lysate and Enriched Phosphopeptides**. The peptides of total cell lysate and enriched phoshopeptides were dissolved in 40 μL 10% NH3·H2O and were separated with self-packed StageTip column (MAGIC C18, 100 Å, 3 μm, Michrom Bioresources^9^) with an elution gradient of 0.5%, 2%, 5%, 8%, 10%, 20%, 40%, 50% and 80%. 8 fractions of each sample were collected in separate tube by centrifuging (800 g) and then combined into 3 fractions (0.5%, 8% and 40% in fraction 1; 2%, 10% and 50% in fraction 2; 5%, 20% and 80% in fraction 3). The samples were lyophilized and stored at -20 °C for further LC-MS/MS analysis.

**Mass Spectrometry-based Proteomics and Phosphoproteomics Analysis**. Dried peptides of each fraction were re-suspended in buffer A (0.1% FA and 0.1% ACN in ddH_2_O). For proteomics, peptides were separated on a home-made capillary column (75 μm i.d. × 15 cm length) packing with 3 μm C18 reverse-phase fused-silica with the EASY-nLC 1000 ultra-performance liquid chromatography (UPLC) system (Thermo Fisher Scientific, San Jose, CA, USA) over a 78 min nonlinear gradient from 6% to 95% acetonitrile at a flow rate of 20 nL/min. For phosphoproteomics, peptides were separated on self-packed capillary column (150 μm i.d. × 12 cm length) packing with 1.9 μm C18 reverse-phase fused-silica with the EASY-nLC 1000 UPLC system over a 78 min gradient from 6% to 95% acetonitrile at a flow rate of 3,000 nL/min. The eluted peptides of total cell lysate and phosphopeptides were ionized and detected by Q-Exactive HF mass spectrometer (Thermo Fisher Scientific, San Jose, CA, USA) and LUMOS mass spectrometer (Thermo Fisher Scientific, San Jose, CA, USA) with a data dependent acquisition (DDA) mode. In brief, for full scan (from m/z 300 to 1400), the spray voltage was set as 2.2 kV and 2 kV, the maximum injection time (MIT) was set to 80 ms and 50 ms, the automatic gain control (AGC) targets were set to 3×10^6^ and 500,000 for proteomics and phosphoproteomics, respectively. The resolution was set to R = 120,000 at m/z 200. For MS/MS scan, the MIT was set to 19 ms and 35 ms, the AGC targets were set to 2×10^4^ and 5,000, the dynamic exclusion for precursor ions was set over a time window of 15 s and 18 s to avoid the repeated detection of the same precursor ions for proteomics and phosphoproteomics, respectively. The resolution was set to R = 15,000 at m/z 200, the isolation width was set to 1.6 m/z and the top ions number was set to 20 with high collision dissociation (HCD) normalized collision energy of 35% which were the same parameters between proteomics and phosphoproteomics.

**Proteomics and Phosphoproteomics Data Analysis by MaxQuant**. Raw data from mass spectrometry of proteomics and phosphoproteomics was parsed by MaxQuant software (version 1.6.0.1) using human database from UniProt (version 201506) with a false discovery rate (FDR) < 0.01 at the level of proteins, peptides and phosphosites. Enzyme specificity was set to trypsin and LysC allowing up to 2 missed cleavages. Fixed modifications were set to carbamidomethyl (cysteine residues). Variable modifications were set to oxidation (M) and acetyl (Protein N-term) for proteomic data, to oxidation (M), acetyl (Protein N-term) and phosphoryl (STY) for phosphoproteomics data. The match-between-runs feature was enabled. The initial mass tolerance of ±20 ppm and final mass tolerance of ±0.5 Da for precursor masses were allowed. The minimum peptide length of 7 amino acids and the minimum andromeda peptide score of 40 were required. A site localization probability of at least 0.75 and a score difference of at least 5 were used as thresholds for the localization of phosphoresidues.

Raw proteomics and phosphoproteomics data for coumarin was obtained from the previous study^1^ where the identical experimental procedure was followed.

**QUALITY CONTROL OF TRANSCRIPTOMICS, PROTEOMICS AND PHOSPHOPROTEOMICS DATA**

**Fold Change Threshold.** To identify the DEGs, DEPs and DEPSs, fold change (FC) thresholds were informed by performing Gaussian fitting analysis and comparing the results for transcriptomics, proteomics and phosphoproteomics (Figure S1). 95% confidence for transcriptomics was determined as: 2SD, FC=2^2SD= 1.268 for 6 h and 1.205 for 24 h for caffeine, FC=1.233 for 6 h and 1.247 for 24 h for coumarin (Figure S1a, b, g, h). 95% confidence for proteomics was determined as: 2SD, FC=2^2SD=1.374 for 10min and 1.381 for 24 h for caffeine, FC=1.360 for 10 min and 1.364 for 24 h for coumarin (Figure S1c, d, i, j). 95% confidence for phosphoproteomics was determined as: 2SD, FC=2^2SD=1.793 for 10 min and 1.560 for 24 h for caffeine, FC=2.263 for 10 min and 1.729 for 24 h for coumarin (Figure S1e, f, k, l). Accordingly, the following FC thresholds were chosen: 1.5 for transcriptomics, 2 for proteomics, and 3 for phosphoproteomics. Therefore, for both caffeine and coumarin, DEGs, DEPs and DEPSs were identified by using a pre-filtering procedure with Benjamin-Hochberg adjusted p-value < 0.05 with the specified FC thresholds.

**Transcriptomics.** The RNAseq experiment carried out on HepG2s exposed to caffeine was able to identify 37,637 features in total (Table S1). The number of detected transcripts was similar across the samples and ranged between 23,000 and 24,900 (Figure S2a). After filtering out the probes with median counts < 5 and normalizing the data with DESeq2^10^ (see Methods), we found that the gene count distribution was kept consistent across all samples (Figure S2b, c). Moreover, a pairwise Pearson correlation coefficient between the samples was high and at the range of 0.94-0.99 (Figure S2d, e).

For the coumarin RNASeq dataset, we were able to identify 58,735 features in total (Table S1). The number of detected transcripts was again similar across the samples and ranged between 22,400 and 25,010 (Figure S3a). Upon removing the genes with median counts < 5 and conducting the normalization with DESeq2^10^ (see Methods), we found that the gene count distribution was consistent across all samples (Figure S3b, c). Similar to the caffeine data, a pairwise Pearson correlation coefficient between the coumarin samples was high and had an average value of 0.99 (Figure S3d, e).

**Proteomics.** The proteomics experiment carried out on HepG2s exposed to caffeine was able to identify 8,179 proteins in total for both of the 10 min and 24 h (Table S1) samples, respectively. Proteins with intensities detected in fewer than 4 out of 21 samples were filtered out. For the remaining proteins, after median normalizationwere conducted, 129 (1.85%) and 160 (2.29%) missing values were imputed, which resulted in 6,987 and 6,988 proteins for the 10 min (Figure S4a) and 24 h (Figure S4b) samples, respectively. The normalized log_10_ intensities of identified proteins ranged from 6 to 12 for both time points (Figure S4c, d) and indicated a high sensitivity in protein identification ^11^. The distribution of protein log_2_ intensities was consistent across the samples (Figure S4c, d). A Pearson pairwise correlation between the samples was high and ranged between 0.88 and 0.96 (Figure S4e, f).

For the coumarin proteomics data, we were able to identify 6,960 and 6,708 proteins in total for the 10 min and 24 h samples (Table S1), respectively. After conducting pre-filtering and median normalization, 109 (1.75%) and 124 (2.07%) missing values were imputed in Perseus software, which resulted in 6,230 and 6,001 proteins for the 10 min (Figure S5a) and 24 h (Figure S5b) samples, respectively. Similar to the caffeine dataset, the normalized log_10_ intensities of identified proteins ranged from 6 to 12 for both time points (Figure S5a, b) and indicated a high sensitivity in protein identification ^11^. The distribution of protein log_2_ intensities was consistent across the samples (Figure S5c, d). Similar to the caffeine data, a Pearson pairwise correlation between the samples was high and ranged from 0.93 to 0.98 (Figure S5e, f).

**Phosphoproteomics.** The phosphoproteomics carried out on HepG2s exposed to caffeine was able to identify 47,236 in total for both 10 min and 24 h samples (Table S1), respectively. The distribution of localization probability showed that the class Ⅲ phosphosites (probability score > 0.75) accounted for 74.95% cases, indicating high confidence in identified phosphosites (Figure S6a). Class I (probability score between 0.25 and 0.5) and class II phosphosites (probability score between 0.5 and 0.75) accounted for only 9.90% and 14.25% were removed from the further analysis (see Methods). Phosphorylated amino acids were classified as: pSer (71.06%), pThr (20.59%), and pTyr (8.35%) (Figure S6a) in proportions similar to the previously reported distribution^12^. Peptides with intensities detected in at least 4 out of 21 titration samples were kept for further analysis. 2,731 (9.87%) and 2,457 (10.35%) missing intensities were imputed following the normal distribution for 10 min and 24 h sample, respectively (see Methods). This resulted in a total of 27,680 and 23,739 phosphosites for the 10 min (Figure S6b) and 24h sample (Figure S6c), respectively. The log_10_ intensities of the phosphosites ranged from 5 to 11 in the 10 min (Figure S6b) and 24 h sample (Figure S6c), which implied a considerable sensitivity of the detection ^11^. Furthermore, the abundance of the phosphosites was found consistent in all of the analysed samples (Figure S6d, e). The pairwise Pearson correlation coefficient of phosphosite intensities between the samples ranged from 0.56 to 0.74 (Figure S6f, g), indicating larger differences between samples treated with different concentrations of caffeine in comparison to the proteomics and transcriptomics data.

For the coumarin phosphoproteomic data, we identified 28,288 and 24,152 phosphosites for the 10 min and 24 h samples (Table S1), respectively. The distribution of localization probability showed similar distribution to the caffeine sample with class Ⅲ phosphosites being dominant and accounted for 70.07% cases, whereas class I and class II phosphosites accounted for only 13% and 16.08%, respectively (Figure S7a). Phosphorylated amino acids were classified as: pSer (75.75%), pThr (20.37%), and pTyr (3.89%) (Figure S7a) in proportions similar to the caffeine treated samples^12^. We applied identical filtering procedure as for the caffeine treated samples (see Methods). 1,324 (8%) and 1,301 (9.19%) missing intensities were imputed for 10 min and 24 h sample, respectively. This resulted in a total of 16,541 and 14,157 phosphosites for the 10 min (Figure S7b) and 24 h sample (Figure S7c), respectively. Considerable sensitivity of the detection was noted with log_10_ intensities of the phosphosites being in the range from 5 to 11 (Figure S7b, c), whereas the abundance of the phosphosites was found consistent after normalization similar to the caffeine treated samples (Figure S7d, e). The pairwise Pearson correlation coefficient of phosphosite intensities between the samples ranged from 0.53 to 0.74 (Figure S7f, g), again confirming larger differences in the coumarin phophoproteomic data in comparison to the proteomics and transcriptomics data.

**ABBREVIATIONS**

EPA, Environmental Protection Agency; NAMs, new approach methodologies; HTTr, High throughput transcriptomics; POD, point of departure; POD_T_/POD_P_/POD_Ph_, POD at the gene/protein/phosphoprotein level; DMEM, Dulbecco’s modified eagle medium; FBS, fetal bovine serum; PBS, Phosphate buffered saline; DEGs, differentially expressed genes; DEPs, differentially expressed proteins; DEPSs, differentially expressed phosphosites; FC, fold change; BMD, benchmark dose; BMDL, the lower bound of BMD; BMDU, the upper bound of BMD; BBMD, Bayesian BMD method; BMR, Benchmark Response; GO, Gene Ontology; MOFA, Multi-Omics Factor Analysis; PPIs, protein-protein interactions; ChEA3, ChIP-X Enrichment Analysis 3; TTN, titin; MLC, myosin light chain; ADORA1, adenosine receptor 1; ADORA2A, adenosine receptor 2A; DTT, Dithiothreitol; IAA, iodoacetamide; FA, formic acid; ACN, acetonitrile; IMAC, immobilized metal ion affinity chromatography; UPLC, ultra-performance liquid chromatography; DDA, data dependent acquisition; MIT, maximum injection time; AGC, automatic gain control; HCD, high collision dissociation; FDR, false discovery rate.
